# Reproductive phasiRNA loci and DICER-LIKE5, but not microRNA loci, diversified in monocotyledonous plants

**DOI:** 10.1101/2020.04.25.061721

**Authors:** Parth Patel, Sandra M. Mathioni, Reza Hammond, Alex E. Harkess, Atul Kakrana, Siwaret Arikit, Ayush Dusia, Blake C. Meyers

## Abstract

In monocots other than maize and rice, the repertoire and diversity of microRNAs (miRNAs) and the populations of phased, secondary, small interfering RNAs (phasiRNAs) are poorly characterized. To remedy this, we sequenced small RNAs from vegetative and dissected inflorescence tissue in 28 phylogenetically diverse monocots and from several early-diverging angiosperm lineages, as well as publicly available data from 10 additional monocot species. We annotated miRNAs, siRNAs and phasiRNAs across the monocot phylogeny, identifying miRNAs apparently lost or gained in the grasses relative to other monocot families, as well as a number of tRNA fragments misannotated as miRNAs. Using our miRNA database cleaned of these misannotations, we identified conservation at the 8^th^, 9^th^, 19^th^ and 3’ end positions that we hypothesize are signatures of selection for processing, targeting, or Argonaute sorting. We show that 21-nt reproductive phasiRNAs are far more numerous in grass genomes than other monocots. Based on sequenced monocot genomes and transcriptomes, DICER-LIKE5 (DCL5), important to 24-nt phasiRNA biogenesis, likely originated via gene duplication before the diversification of the grasses. This curated database of phylogenetically diverse monocot miRNAs, siRNAs, and phasiRNAs represents a large collection of data that should facilitate continued exploration of small RNA diversification in flowering plants.

## Introduction

Small RNAs (sRNAs) are key regulators of gene expression at the transcriptional and post-transcriptional level. MicroRNAs (miRNAs), a class of small non-coding RNAs with lengths ranging from 20 to 22 nucleotides (nt), are generated from stem-loop precursor RNAs processed by the RNase III family enzyme DICER-LIKE1 (DCL1), which yields a miRNA/miRNA* duplex. The duplex has 2-nt overhangs in the 3’ ends; these ends are methylated by the methyltransferase HUA ENHANCER1 (HEN1) for protection from degradation (Johnson et al., 2009; Yang et al., 2006). Generally, one strand of the duplex, the miRNA, is loaded into an Argonaute protein (AGO1) to form the RNA-induced silencing complex (RISC). RISC, via sequence homology to the miRNA, recognizes target messenger RNAs (mRNAs), and this interaction causes post-transcriptional repression by either target mRNA cleavage or translational repression (Axtell, 2013). miRNAs are involved in a multitude of plant biological processes such as seed germination (Reyes and Chua, 2007; Liu et al., 2007), leaf morphogenesis (Palatnik et al., 2003), floral development (Mallory et al., 2004), and responses to biotic (Li et al., 2011; Zhang et al., 2016) and abiotic stresses (Leung and Sharp, 2010; May et al., 2013).

PhasiRNAs, another important sRNA class, are distinguished from miRNAs in their biogenesis. PhasiRNA biogenesis starts from a single-stranded product of RNA Polymerase II (Pol II) derived from a genomic *PHAS* locus that is capped and polyadenylated as a typical mRNA. Next, cleavage of this mRNA is directed by a 22-nt miRNA (Zhai et al., 2015; Cuperus et al., 2010). The cleaved phasiRNA precursor is made double stranded by RNA-DEPENDENT RNA POLYMERASE 6 (RDR6), and then this double-stranded RNA (dsRNA) is processed in an iterative or “phased” manner (i.e. into consecutive 21- or 24-nt sRNAs) by a Dicer-like enzyme. DCL4 and DCL5 produce 21- and 24-nt phasiRNAs, respectively (Song et al., 2012). The 21-nt phasiRNAs are widespread in plants, originating in land plants hundreds of millions of years ago as the *trans*-acting siRNA (tasiRNA) loci, but also derived from diverse protein-coding gene families (Fei et al., 2013; Xia et al., 2017). The 24-nt “reproductive” phasiRNAs have been described only in angiosperms, are highly enriched in anthers, and are typically but not always triggered by miR2275 (Kakrana et al., 2018; Xia et al., 2019). A special class of 21-nt “reproductive” phasiRNAs are highly expressed in pre-meiotic anthers of some monocots, triggered by miR2118 (Johnson et al., 2009; Kakrana et al., 2018). The function of either class of phasiRNAs is still not clear but perturbations of both result in defects in male fertility (Fan et al., 2016; Komiya et al., 2014; Ono et al., 2018; Zhang et al., 2018). miRNAs have been investigated in many plant species, both in individual genomes and from limited-scale comparative analyses (You et al., 2017; Montes et al., 2014).

In the monocots, a group of about 60,000 species, most studies of sRNAs have focused on members of the Poaceae (grasses), with scant data from non-grass monocots (Kakrana et al., 2018). Rice, *Brachypodium,* and maize are the most studied of the grasses, with miRNAs characterized using varying genotypes, tissue types, growth and stress conditions (Jeong et al., 2011; Zhang et al., 2009). With the major goal of assessing the diversity and origins of miRNAs in monocots, we analyzed sRNA data from 38 phylogenetically diverse monocots, spanning orders from the Acorales to the Zingiberales. We described sRNA size classes, miRNA conservation, divergence, sequence variability, 5’ and 3’ end nucleotide preferences, and single nucleotide sequence profile characterizing positional biases and providing novel insights within the miRNA sequences. We performed comparative analysis of miR2118 and miR2275 and their long non-coding RNA (lncRNA) targets in monocots relative to other flowering plants, demonstrating their presence and absence in these species. We found that both miR2118 and miR2275 are conserved across diverse monocot species, and are present in vegetative tissues but are found at high abundances predominantly in inflorescence tissues. The 21- and 24-nt *PHAS* loci are most numerous in the genomes of grasses, relative to other monocots, and are similarly most abundant in inflorescence tissues. Fewer *PHAS* loci were identified in non-grass monocots. Overall, our study provides a deep comparative analysis of sRNAs in monocots, including a refined database of monocot miRNAs.

## Results

### Sequencing from diverse monocots demonstrates atypically abundant 22-nt siRNAs

We collected materials and sequenced sRNAs from 28 monocot species spanning nine taxonomic orders: Poales (17 species), Arecales (3 species), Zingiberales (2 species), Commelinales (1 species), Asparagales (6 species), Pandanales (1 species), Liliales (1 species), Alismatales (6 species), and Acorales (1 species) (Supplemental Table S1). These species included an early-diverging monocot *Acorus calamus* (Acorales) and the early diverging Poales (grasses) *Pharus parvifolius*, *Anomochloa marantoidea*, and *Streptochaeta angustifolia* (Kellogg, 2001). The sRNA libraries from these species totaled 52 vegetative and 148 inflorescence or reproductive tissues; these 200 sRNA libraries yielded 5,312,866,505 total sRNA sequences after trimming and quality control of reads (Supplemental Table S2). For some analyses, we utilized public sRNA data from an additional ten monocot species (Figure 1; Supplemental Table S1). We also included sRNAs from *Nymphaea colorata* (Nymphaeales), *Amborella trichopoda* (Amborellales), and *Arabidopsis thaliana* as outgroups (Figure 1). In total, our study comprised billions of sRNAs from 41 diverse angiosperm species.

**Figure 1.**
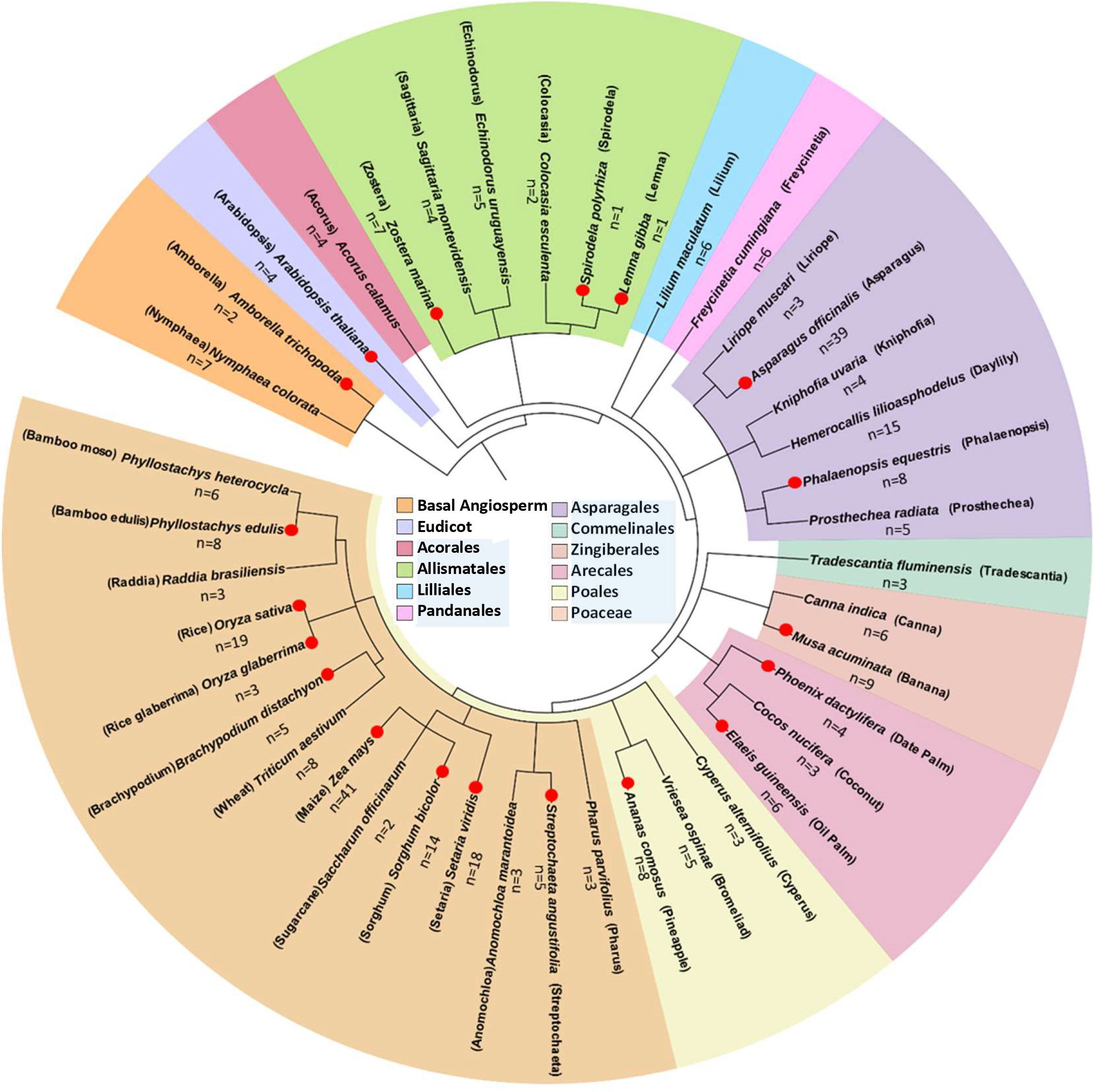
Phylogenetic distribution of species sampled for small RNAs. *N. colorata*, *A. thaliana* and *A. trichopoda* were included for comparative purposes. Orders of the monocots are shaded in light blue in the legend. Red dots denote species with genome sequences available at the time of this work. The phylogeny was generated using phyloT (phylot.biobyte.de) based on NCBI taxonomy. This phylogenetic tree was annotated using iTOL (https://itol.embl.de).

We first assessed variation in the size distribution of small RNAs across the monocots. After removing reads <18 nt and >36 nt, 21 and 24 nt long sRNAs are the two dominant sizes (Figure 2, Supplemental Figure S1, Supplemental Table S2). These 21- and 24-nt peaks are typically comprised of miRNAs (21 nt) and heterochromatic siRNAs (hc-siRNAs; 24 nt), but in the case of anthers may also include a large number of abundant phasiRNAs (Zhai et al., 2015; Johnson et al., 2009). The Nymphaeales and Asparagales displayed the highest relative proportions of 21- and 24-nt sRNAs compared to other taxonomic orders, respectively (Figure 2A). For the majority of sampled species, the prominent sRNA peak was 24 nt, except for the Nymphaeales and Pandanales in which the 21 nt peak was more prominent. In the Zingiberales, 21 and 24 nt size classes are similarly abundant.

**Figure 2.**
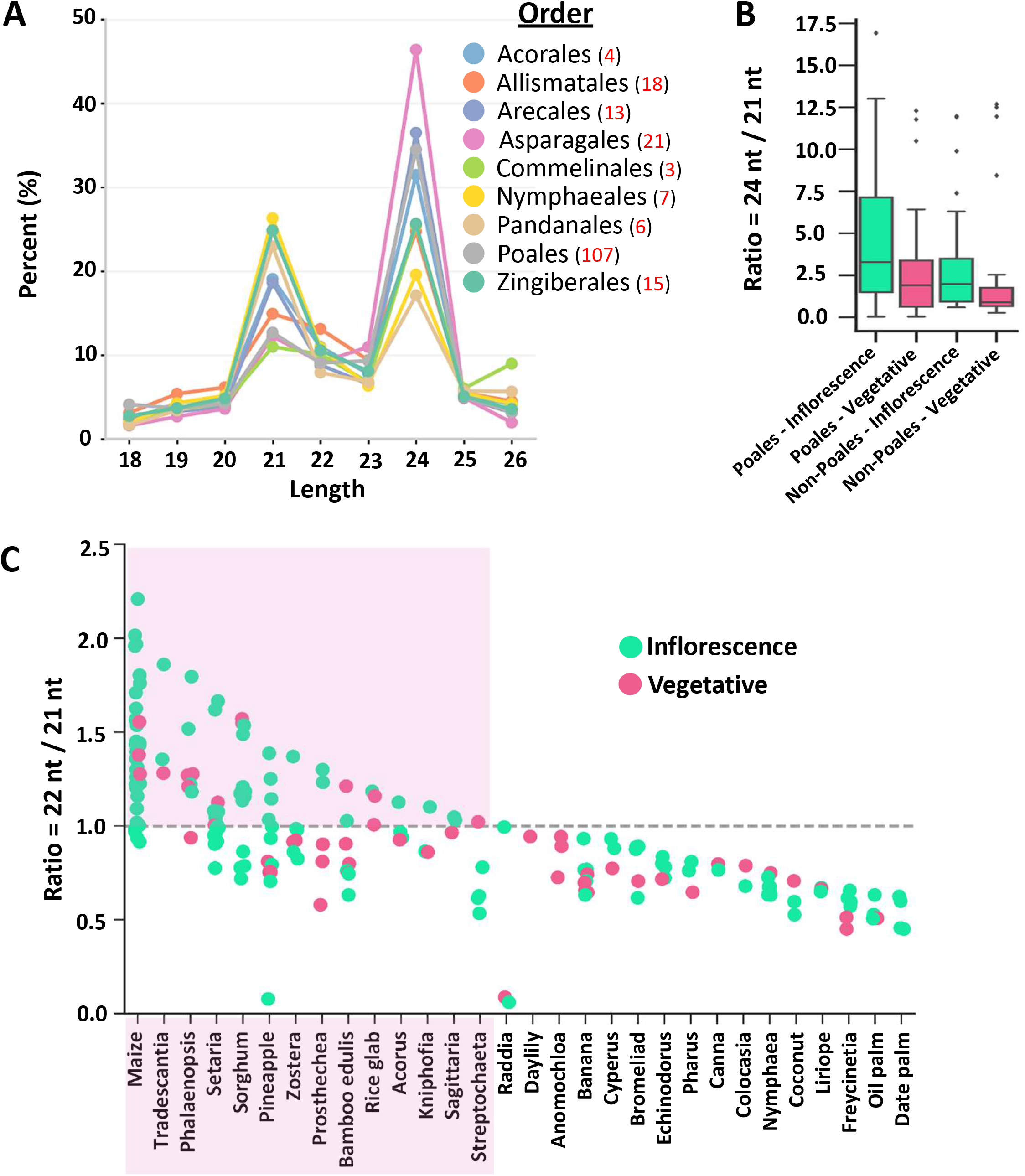
Small RNA size distribution variation across the monocots and their tissues. (A) The relative proportion of sRNAs was calculated as percentages (Y-axis) for each size category (X-axis) in total reads in 200 sequenced libraries across 28 plant species grouped in nine different plant orders along with number of libraries in that order, denoted by red font. Reads longer than 26 nt were not included in this study. (B) Box plots compare the ratio of the 24- and 21-nt sRNA reads distribution (Y-axis) between the Poales (grasses) and non-Poales, defined as all species except the grasses (X-axis). The center line (black line) in each plot indicates median of the distribution. (C) Dot plot depicts the ratio of the 22- and 21-nt distinct sRNA reads (Y-axis) among species (X-axis), grouped by vegetative (pink dots) and Inflorescence libraries (green dots). Highlighted pink box indicates species for which the ratio is ≥ 1. Dotted grey line denotes equal ratio (value of ‘1’ on Y-axis).

To complement the global sRNA abundance analysis, we next calculated the ratio of the distinct (unique) sequences of 24 nt to 21 nt sRNAs; since grasses have been well characterized, we compared species in the Poales (grasses) to non-Poales (all monocot data in our study except the grasses), and inflorescence versus vegetative tissues (Figure 2B). Overall, the Poales display a higher proportion of 24 nt sRNAs than non-Poales across all libraries, perhaps indicative of more 24 nt hc-siRNAs or 24 nt phasiRNAs.

Next, we identified species with a disproportionately high level of 22 nt sRNAs, as our prior work has identified an unusual, RDR2-independent class of 22-nt sRNAs in maize (Nobuta et al., 2008). Our recent work using machine learning approaches demonstrated that these maize 22-nt siRNAs have distinct sequence characteristics (Patel et al., 2018). Yet, outside of these reports in maize, there has been a paucity of data on these 22-nt sRNAs in the last decade, perhaps because monocot small RNAs are so poorly characterized. We computed the ratio of the distinct 22 and 21 nt sRNAs, again comparing inflorescence and vegetative tissues (Figure 2C). This ratio exceeded 1 (i.e. higher levels of 22-nt sRNAs) for most grasses (*Setaria*, *Sorghum*, etc.) and several non-grass monocots (*Tradescantia*, *Phalaenopsis*, *Zostera*, etc.) (Figure 2C). This higher proportion of 22-mers was also more often observed in inflorescence than vegetative tissues. This result is consistent with a widespread occurrence of these poorly characterized 22 nt siRNAs in monocot species other than in just maize; their biogenesis and roles are yet to be described.

### Identification of miRNAs variably conserved within the monocots

With these data, we sought to characterize miRNAs present in monocots, and those that either pre-date the split with eudicots or likely emerged since then. Since validated miRNAs longer than 22 nt are rare, our analysis focused on the 20, 21, and 22 nt lengths. We utilized two main strategies for the miRNA analysis. First, we identified conserved candidate miRNAs using a custom, homology-based pipeline, using mature plant miRNAs from miRBase to query all the sRNAs (see Methods for details). This analysis yielded 84,390 miRNA sequences; these were filtered to find those with an abundance of ≥100 reads in at least one sequencing library, and yielding 5,354 miRNA sequences. These sequences belonged to 290 distinct miRNA families (all annotated in miRBase); the number of miRNA families per species is shown in Supplemental Table S3. We separated these miRNAs into those that are highly conserved, intermediately conserved, and not conserved – categories described in the following paragraphs.

#### Conserved miRNAs

We identified six highly conserved miRNAs found in more than 34 species; these are the well-known miRNAs miR156, miR165/166, miR167, miR171, miR319, and miR396 (Figure 3A; Supplemental Table S4, Part A). A set of another fifteen well-conserved miRNAs were found, present in 20 to 34 species (Figure 3A; Supplemental Table S4, Part A). Another set of miRNA families were observed in 10 to 20 of the 41 species, which for descriptive purposes we state as having a moderate or intermediate level of conservation (Figure 3B; Supplemental Table S4, Part B). The absence of a miRNA family from one species does not imply that it is not encoded in that genome as some miRNAs are tissue-specific and our sampling and depth of sequencing was not an exhaustive analysis. However, these data were useful as a representative set of monocot miRNAs.

**Figure 3.**
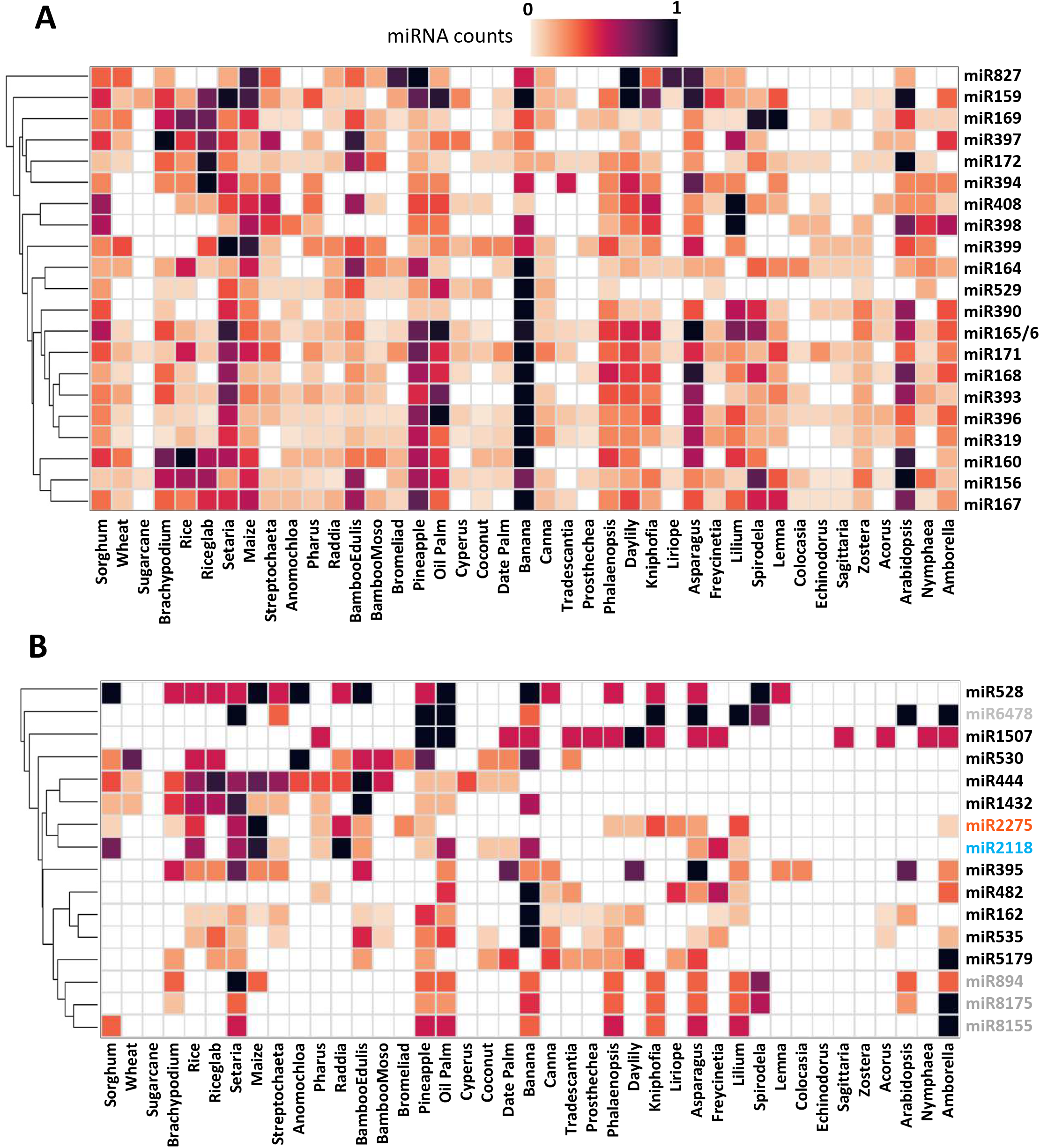
Variable levels of conservation in monocot-specific miRNAs. miRNA abundances were assessed using the sRNA data from vegetative and reproductive tissues. A sequence with >= 100 reads in either vegetative or inflorescence tissues was retained. miRNA families were divided into two groups according to their conservation, defined as highly conserved and intermediate conserved. Heatmap colors represent row normalized miRNA candidates counts in the family ranging from 0 (white) to 1 (black) as illustrated in the color keys. The miRNAs were hierarchically clustered based on counts using “single” method and “euclidean” distance. (A) High conservation was defined by miRNA families identified in more than 19 species out of all 41 species examined. (B) Intermediate conservation was defined by miRNA families identified between 10 and 19 species. miR2118 and miR2275, the reproductive phasiRNA triggers in the grasses, are denoted by blue and orange fonts, respectively. miR6478, miR894, miR8155 and mi8175 are in light grey as all correspond to tRNA fragments (see main text).

We made a number of observations about monocot miRNAs. First, most “three-digit miRNAs” (miRNAs with numbers lower than miR1000, i.e. miR166, miR399, miR827, etc.) are conserved across monocots (Figure 3A). Second, the data from banana showed a strong pattern for higher counts of miRNA candidates for both highly conserved (Figure 3A) and intermediately conserved (Figure 3B) groups. Third, *Colocasia*, *Echinodorus*, *Sagittaria*, and *Zostera* show a poor representation of highly conserved miRNAs, and essentially no moderately conserved miRNAs, except for miR395 in *Colocasia* and miR1507 in *Sagittaria*. And finally, sugarcane showed weak or no representation of conserved miRNAs. The poor result in sugarcane could be attributed to the sampled tissue type, technical complication due to a highly polyploid genome, an issue of sample preparation, or reads not exceeding the abundance of 100 in the two libraries used in this analysis.

We also noted several intriguing patterns of miRNA conservation in the monocots (Fig. 3B). First, miR1507, a trigger for 21-nt phasiRNAs from nucleotide binding and leucine-rich repeat (NB-LRR) pathogen-defense genes in legumes (Fei et al., 2015) is not present in the grasses (except in *Pharus*), but it is present in other monocots, perhaps indicative of a lineage-specific loss. Second, miR444, miR530, and miR1432 showed strong representation in grasses and poor representation outside of the grasses, perhaps consistent with recent evolutionary emergence. Third, miR482 is detected in early-diverged monocots but not in the grasses, consistent with earlier reports of functional diversification of miR482 and miR2118 (Xia et al., 2015). Fourth, miR894, miR8155, and miR8175 showed a similar pattern of presence across the sampled monocots, suggesting these miRNAs may comprise a family. Fifth, we identified several miRNAs present specifically as 22-mers in *Amborella*, including miR482, miR1507, and miR2275; their presence in the sister to all flowering plants suggests these miRNAs emerged prior to the monocots.

We examined several of these observations in more detail, starting with the set of three miRNA families (miR894, miR8155, and miR8175) demonstrating similar patterns of representation (Figure 3B). These miRNAs have been separately described in eudicots, mosses and other lineages (Montes et al., 2014; Harkess et al., 2017), but not previously been shown to have a common origin. We aligned the family members and found a high degree of similarity that is suggestive of a superfamily (Supplemental Figure S4A). A review of the literature mentioning these three miRNAs found that miR894 was previously inferred to be a tRNA fragment (Montes et al., 2014). Therefore, we analyzed miR8155 and miR8175 to determine if these might also be tRNA fragments. The annotated copy of the Arabidopsis miR8175 corresponds to the 3’ end of tRNA Asp-GTC-8-1 (Supplemental Figure S4B); while miR8155 from oil palm corresponds to the 3’ end of annotated tRNA from *Oryza sativa* (orysa-Met_iCAT-53) (Supplemental Figure S4C). These misannotations may be perpetuating confusion about these small RNAs and thus the miRNAs should be blacklisted or scrubbed from databases.

We next examined the set of monocot-specific miRNAs represented by miR444, miR530, and miR1432. miR444 has been previously characterized in several grass species (Lu et al., 2008), and it was identified without further analysis in pineapple (Md Yusuf et al., 2015). We found miR444 in several other monocots including the palms, but no earlier diverging lineages (**Figure 3B**). A characteristic of miR444 in grasses is its genomic antisense configuration relative to the target gene (Lu et al., 2008); we observed the same configuration in pineapple, indicating that this arrangement may reflect its ancestral state and even its evolutionary origins (Supplemental Figure 7). miR530 and miR1432 were also described previously in pineapple (Md Yusuf et al., 2015), and we also found that they were detected in other sister species within the commelinids, but not earlier in the monocots. Therefore, our analysis of annotated miRNAs demonstrated a combination of patterns of conservation and divergence within the monocots, with at least three monocot-specific miRNAs that emerged coincident with the commelinids.

#### Novel miRNAs predicted as conserved within monocots

Next, we tested if our data revealed any monocot-wide novel miRNAs that are not annotated in miRBase. We utilized *de novo* miRNA prediction for the monocot species for which a genome sequence is available at the time of this analysis (15 species, in 2018) and we compared these across the sRNA data of all analyzed species. This identified novel and weakly conserved miRNA families. For the 15 species with genomes, using small RNA data from this study and publicly available data, we used a new pipeline called *miRador* (available on Github, see methods) and cross-checked the results using the well-established ShortStack pipeline (Johnson et al. 2016). Both pipelines implement the strict, recently-described criteria for miRNA annotation (Axtell & Meyers, 2018). In those criteria, predicted miRNAs with five or fewer nucleotide differences were then classified as members of a single miRNA family. We did not consider candidates found in only one genome, as these are harder to validate in a large-scale screening, and are thus prone to misannotation. The result of both pipelines was similar, with no candidates for novel miRNAs conserved across all 15 monocot species. There was one case for which novel miRNAs appeared to be conserved across at least two species, *Setaria* and *Sorghum* (Supplemental Table S5). These miRNAs passed our strict annotation criteria, and target prediction plus analysis of PARE data generated for this study from multiple tissues (Supplemental Table S2) found no validated targets in either genome, so their possible functions or roles in post-transcriptional silencing remain unclear. Overall, there is scant evidence for the presence of any monocot-wide, conserved and novel miRNAs.

### Size distribution of conserved miRNAs displays strong conservation of length in monocots

Plant miRNAs are typically 21- or 22-nt, with this length difference often determining whether or not they function to trigger phasiRNAs; therefore, we were interested to determine whether conserved miRNAs are also conserved in their length. We computed the distribution of miRNA sizes as a percentage (between 0 and 100) of the total abundance for highly and intermediately conserved miRNA families (Figure 4). In the 37 miRNA families (Figure 4A, B), 21 nt was the most frequent length. Four miRNA families (miR156, miR394, miR395 and miR6478) exhibited substantial proportions of 20-nt sequences (>30% of abundance). This length for both miR156 and miR394 is due to formation of asymmetric bulges and mismatches in the duplex (Lee et al., 2015). We were curious about miR6478, predominantly a 20 nt miRNA across multiple species in our data, yet reported previously only in poplar and rice (He et al., 2015; Puzey et al., 2012). We observed that it was present in Arabidopsis and *Amborella,* but not annotated in either species. When we analyzed the sequence in the Arabidopsis genome (Supplemental Figure S8), it corresponds to tRNA precursors, and thus we infer that this is a tRNA fragment (tRF) and not a true miRNA. This matches with prior conclusions about a different miRNA included in our work, miR894, also inferred to be a tRNA fragment (see above) (Montes et al., 2014).

**Figure 4.**
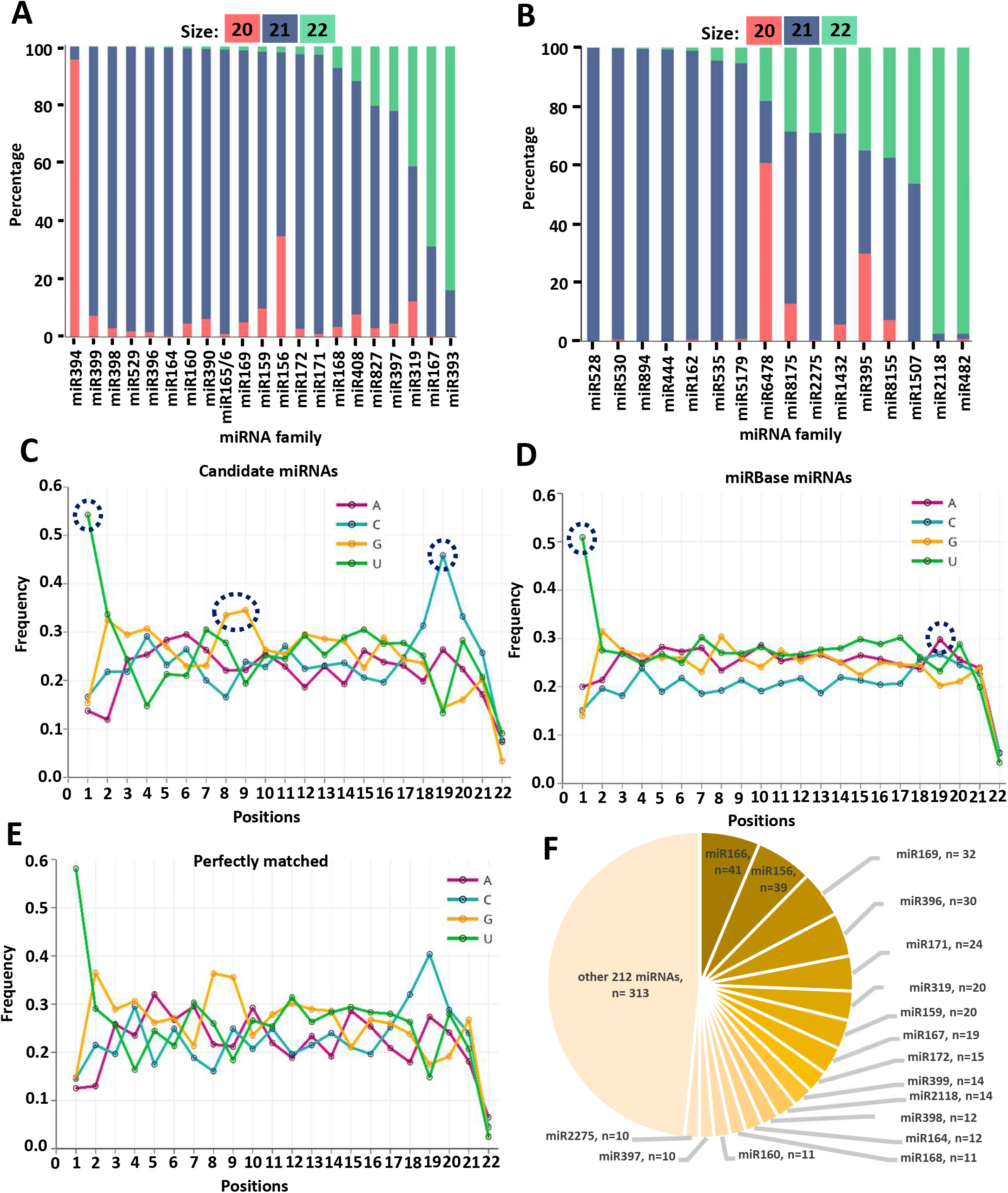
Size distribution and sequence profile characterize position-specific nucleotide biases of conserved miRNA variants. (A-B) The stacked bar plots show the size distribution of reads as a percentage (between 0 and 100) of abundance out of total abundance (Y-axis, denoted 20-mers in red; 21-mers in blue; 22-mers in green) in the conserved miRNA families. As in Figure 3, conservation was defined by miRNA families identified in more than 10 species out of all 41 species examined. Bar plots are sorted from low to high percentage of 22-mers; abundances were combined for all species in which the miRNAs were detected. (A) Size distribution in the most conserved 21 miRNA families (X-axis). (B) Size distribution in the intermediate conserved 16 miRNA families. (C, D) Single-nucleotide sequence profiles of unique candidate miRNAs (n=2304) (panel C) and unique mature miRNA sequences (n= 3722, size 20 to 22) from miRBase, version 21 (panel D). The frequencies of each of the four bases (A, C, G, and U) at each position are indicated as an open circle. (E) Single-nucleotide sequence profiles of unique candidate miRNAs from panel C perfectly matching (n= 647, panel E) to the miRBase miRNAs. (F) Pie chart illustrating counts of miRNA candidates 100 percent identical to miRBase miRNAs from panel E.

Five miRNA families (miR167, miR393, miR482, miR1507, and miR2118) preferentially accumulated as 22 nt sRNAs. Length variation for miR167 and miR393 was observed previously, although miR393 was reportedly rarely as 22 nt in monocots, perhaps reflecting a narrow set of sampled species in this lineage (Montes et al., 2014). The 22-nt size specificity among miR482, miR1507, and miR2118, reflects well-described roles as triggers of phasiRNAs. miR2275, also a well-known trigger of phasiRNAs (Johnson et al., 2009) was, perhaps unexpectedly, not among the set of preferentially 22 nt miRNAs. The most highly abundant 21 nt sequence (zma-miR2275b-5p) turned out to be a miRNA* (“microRNA-star”) sequence from these loci; this sequence has no known function or targets, so its extraordinary accumulation may reflect something unusual about miR2275, which is the only miRNA reported to date to trigger 24-nt phasiRNAs.

### Single-nucleotide miRNA sequence profiles characterize position-specific nucleotide biases of conserved miRNA variants

Next, we characterized candidate miRNA sequences in greater detail, at the single nucleotide level. We first generated a non-redundant set of 2,304 candidate miRNA sequences from the set of 5,354 sequences of conserved miRNAs (in 290 distinct miRBase-annotated families) found across our libraries. We computed single-nucleotide sequence profiles for these sequences, determining the frequencies of each nucleotide (A, C, G, and U) at each position (Figure 4C). Combining these results, we made several observations: (1) in miRNA candidates, there was a 5’ nucleotide preference for U, consistent with prior reports (Montes et al., 2014; You et al., 2017); (2) a peak of G was observed at the 8th and 9th positions; (3) in the 3’ end of the candidate miRNAs, we observed a peak of C at the 19th position (with a depletion of G and U). For comparison, we plotted the sequence profile of 3,722 distinct mature miRNAs directly from miRBase (version 21; size 20 to 22 nt). Several aforementioned features were conserved, except for the peak of G at the 9th and C at the 19th positions, and the miRBase miRNA sequence profile lacked distinctive sequence characteristics (Figure 4C, 4D). To understand the basis of this difference, we assessed the profile of the subset of our 2,304 miRNAs used for Figure 4C that perfectly match miRBase-annotated miRNAs. This yielded two lists: (1) unique candidate miRNAs from panel C perfectly matching miRBase (n=647, analyzed in Figure 4E), and (2) those not perfectly matching miRBase (n=1656, Supplemental Figure S3). The majority of these perfectly-matched candidate miRNA sequences are well known miRNAs (Figure 4F). We observed similar sequence profiles of candidate miRNAs whether or not they perfectly matched to miRBase miRNAs; since the miRBase miRNAs lacked these distinctive signals, there may be “contaminating” annotations among miRBase miRNA that dilute the signal. This is supported by a recent commentary (Axtell and Meyers, 2018) which suggested that miRBase in its current state contains many low confidence and erroneous annotations of miRNAs.

We next characterized and investigated the previously unreported signatures in these plant miRNAs that we observed at the 8th, 9th, and 19th positions, and test if these observations are consistent across eudicots as well. We focused on Arabidopsis miRNAs from miRBase, filtering them based on their abundance in publicly available sRNA expression data (from GEO series GSE44622, GSE40044, GSE61362, and GSE97917). We used a normalized abundance cutoff of 1000 TP2M (1,000 transcripts per 2 million mapped reads) and we retained distinct sequences of size between 20 and 22, rendering a total of 138 sequences. Among these sequences, we observed the conserved pattern of G at 8th and 9th positions, but a peak of A rather than C at the 19th position (Supplemental Figure S6A). To assess the nature of this A at the 19th position (“19A”), we segregated all the miRNAs by their 5’ nucleotide, creating four more plots. The 5’ U miRNAs, the 5’ end typical of miRNAs (Mi et al., 2008), uniquely displayed a 19C (Supplemental Figure S6B), whereas the other three classes had 19A (Supplemental Figure S6C to S6E). This 19C could be evolutionarily advantageous for 5’ U miRNAs because in a 21-nt miRNA this would yield a 5’ G on the complementary strand, i.e. the “passenger” or miRNA* strand. Since 5’U is a strongly favored nucleotide across the majority of miRNA families and few mature miRNAs have 5’ G, this nucleotide composition may contribute to AGO sorting, loading, or binding. An alternative hypothesis is that many 5’G/19A miRNAs may be mis-annotated passenger strands.

### 5’ and 3’ nucleotide features of conserved miRNAs

Because the 5’ nucleotide of miRNAs is a distinguishing feature, mainly for AGO sorting and hence function (Mi et al., 2008), we analyzed the 5’ and 3’ ends of the sRNAs described above. We characterized the 5’ nucleotide prevalence in the 21 highly conserved and in the 16 intermediately conserved miRNA families, and focused on miRNAs from 20 to 22 nt. In the majority of miRNA families, U was the most prevalent 5’ nucleotide, consistent with earlier reports (Montes et al. 2013 and You et al. 2017) (Figure 5A, 5B). We observed several exceptions at the 5’ end: miR390, miR529, and miR172 predominantly displayed an A at the 5’ position, consistent with earlier reports (Montes et al. 2013 and You et al. 2017). In the intermediately conserved miRNA families, a 5’ U also predominated (Figure 5B). The exceptions we found (miR6478 and miR894 with a 5’ C, and miR8155 and miR8175, with a uniform distribution of G, C, and U), were all tRNA fragments (see above), suggesting that these 5’ nucleotides can support the segregation of high quality versus suspicious miRNA annotations.

**Figure 5.**
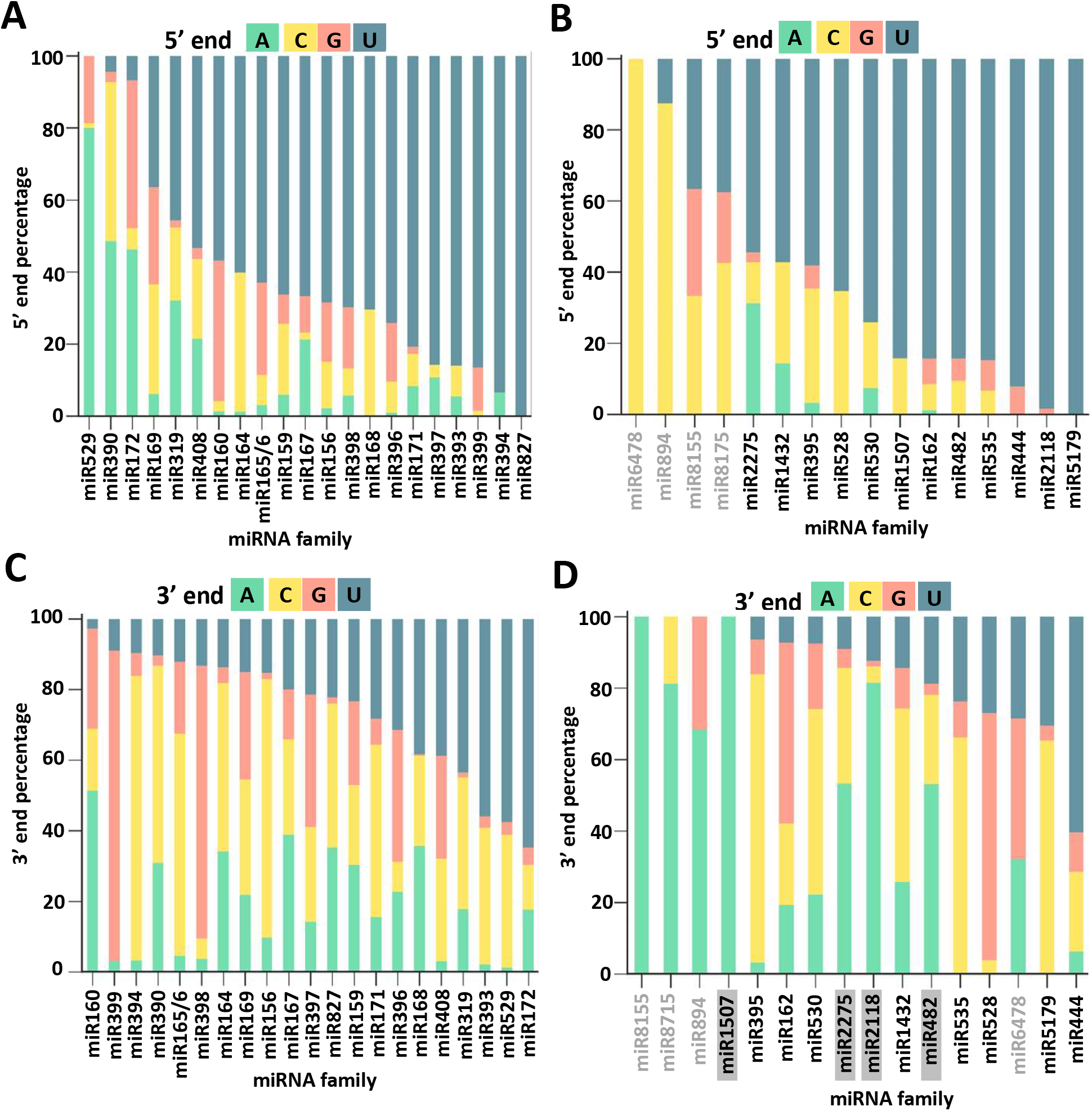
5’ and 3’ nucleotide distribution of conserved miRNA families. The stacked bar plots show the 5’- and 3’-end nucleotide composition as a percentage (between 0 and 100) of all four nucleotides in the conserved miRNA families. Conservation is defined by miRNA families identified in more than 10 species out of all 41 species examined. Bar plots are sorted from low U to high U percentage. (A) The 5’ nucleotide composition (Y-axis) in the 21 most-conserved miRNA families (X-axis). (B) The 5’ nucleotide composition in the 16 intermediate-conserved miRNA families, with the misannotated tRNA fragments noted in grey text, all enriched in 5’ C or G. (C) The 3’ nucleotide composition in the 21 most-conserved miRNA families. (D) The 3’ nucleotide composition in the 16 intermediate-conserved miRNA families. The four known triggers of phasiRNAs are highlighted with grey boxes while the tRNA fragments are noted with grey text.

The 3’ terminal nucleotide identity of miRNAs is not well studied, although we have previously reported that this position may be more important than the seed region (positions 2 to 13), for miRNA-target interactions generating secondary siRNAs (Fei et al., 2015). Hence, we examined the 3’ nucleotide prevalence in the highly and intermediately conserved miRNA families. We observed a 3’ C in the majority of miRNA families (Figure 5C, 5D). A 3’ U was the second most-prevalent nucleotide (Figure 5C, 5D). The 3’ nucleotide identity in a few miRNA families was different. For example, over half of the miR398, miR399, and miR528 family members were enriched in 3’ G (Figure 5C, 5D). The tRNA fragments miR894, miR8155, and miR8175 again had terminal nucleotides inconsistent with “true” miRNAs, in this case with high proportions of 3’ A. The four 22-nt miRNA family members, known triggers of phasiRNAs (miR482, miR1507, miR2118, and miR2275) were all enriched for A in the 3’ end, a 3’-end bias perhaps required for target interactions that instigate the biogenesis of phasiRNAs (Figure 5D). Thus, 3’ nucleotide conservation in plant miRNA families may be as important for miRNA function as the 5’ end, with special importance for triggers of phasiRNAs.

### The biogenesis pathway for reproductive phasiRNAs diversified in the monocots

We next focused on a pathway that has been well described in monocots, the reproductive phasiRNAs. These phasiRNAs are enriched in anthers and require specialized miRNA triggers and long non-coding RNA precursors, as well as the monocot-specific DICER-LIKE 5 (DCL5) protein (Kakrana et al., 2018; Zhai et al., 2015; Xia et al., 2019).

### miR2118 and 21-*PHAS* loci

The trigger of 21-nt phasiRNAs, miR2118, was detected in 16 of the 41 species surveyed in this study (Figure 6). This miRNA is highly abundant (>100 reads) in vegetative tissues of *Raddia*, oil palm, banana, asparagus, and *Freycinetia*; in vegetative tissues of eudicots, *NLR* disease resistance genes are the most common target of the miR2118 superfamily (Zhai et al., 2011). Read abundances below 100 were observed in other monocot species, not only in the Poales but also in the Alismatales, and also in *Amborella trichopoda*. In the inflorescence tissues, miR2118 was highly abundant in most of the grasses, in the order Arecaceae, and in several other monocots (banana, asparagus, and *Freycinetia*). *Nymphaea colorata* (Nymphaea) and *Amborella* also have miR2118 with read abundances lower than 100 (Figure 6). These results confirmed the widespread nature of miR2118 in angiosperms, with high abundance predominantly in the inflorescence tissues of monocot species outside of the grasses.

**Figure 6.**
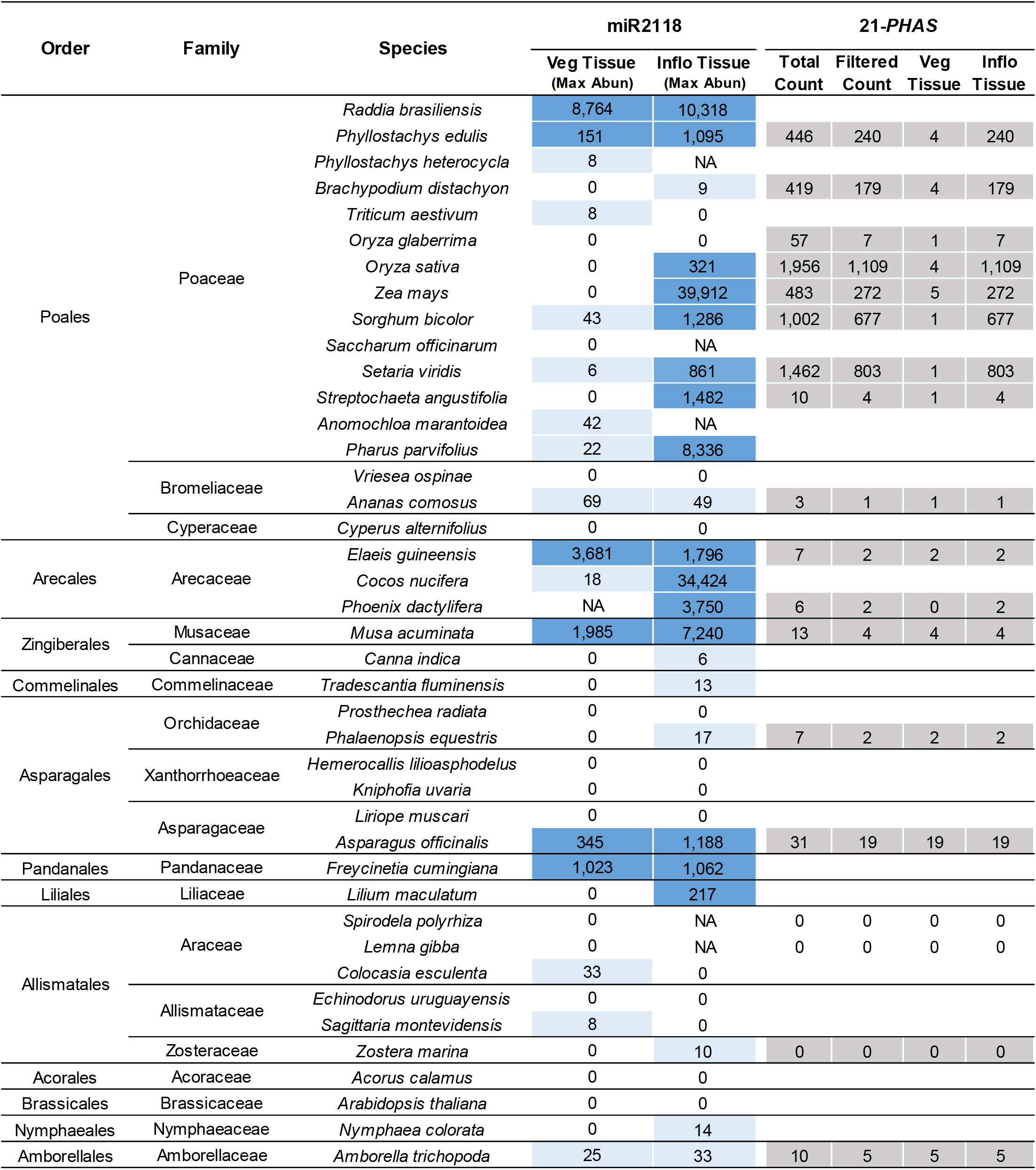
miR2118 abundance and 21-*PHAS* counts in monocots. The miR2118 maximum abundance is shown for each monocot tissue and species where it was identified. The total and filtered 21-*PHAS* counts is shown for each monocot tissue and species where it was identified. The filtered 21-*PHAS* counts for vegetative (Veg) and inflorescence (Inflo) tissues are also shown. Color legend for miR2118 abundance: light blue < 100 reads, dark blue ≥ 100 reads. NA= not available.

Next, we ran a phasing analysis to identify 21-nt phasiRNA-generating loci (21-*PHAS* loci) for 18 species with an available genome sequence, yielding numerous 21-*PHAS* loci (Figure 6). Grasses showed the highest counts of 21-*PHAS* loci, which were more abundant in the inflorescence tissues, with few loci showing abundance in the vegetative tissues – these were typically *TAS3,* important for land plant development (Xia et al., 2017). This high count of 21-*PHAS* loci in grass inflorescence tissues is consistent with their functional importance for anther fertility, for instance as reported in rice (Fan et al., 2016). Outside of the grasses, fewer 21-*PHAS* loci were identified and these showed similar abundances in both inflorescence and vegetative tissues, consistent with TAS3, as observed in pineapple, oil palm, date palm, banana, *Phalaenopsis*, asparagus, and *Amborella* (Figure 6).

We also performed *de novo* trigger prediction to characterize the miRNA triggers of 21-*PHAS* loci, to help determine their potential roles and biogenesis. In the grasses, the biogenesis of reproductive phasiRNAs is typically dependent on one of two 22-nt miRNA triggers (miR2118 and miR2275, for 21- and 24-nt phasiRNAs), while *TAS3* is triggered by miR390. The majority of the 21-*PHAS* loci in the grasses are triggered by miR2118 (Table 1). In the monocots outside of grasses, there were comparatively few 21-*PHAS* loci and only a subset of these had miR2118 as trigger, indicating other miRNAs may function as triggers; this is not unexpected as there are many miRNAs that trigger 21-nt phasiRNAs from diverse targets (Fei et al., 2013). In fact, a proportion of these were *TAS3* (Table 1). However, the relative absence of 21-nt reproductive phasiRNAs outside of the grasses suggests that they are less prevalent, although it is possible that we did not sample the correct anther stage.

**Table 1.** 21- and 24-*PHAS* loci with miRNA triggers for sampled genera. Species are listed only for which a genome was available, and a single representative for each genus is shown. The number of 21-*PHAS* loci that are *TAS3* loci are denoted in parenthesis.

| Order | Family | Species | Total Count of 21-*PHAS* loci | No. of loci with miR2118 as a trigger | Total Count of 24-*PHAS* loci | No. of loci with miR2275 as a trigger |
| --- | --- | --- | --- | --- | --- | --- |
| Poales | Poaceae | *Phyllostachys edulis* | 446 (4) | 373 | 25 | 16 |
|  |  | *Brachypodium distachyon* | 419 (2) | 362 | 217 | 186 |
|  |  | *Oryza sativa* | 1956 (9) | 1810 | 126 | 101 |
|  |  | *Zea mays* | 483 (6) | 395 | 246 | 134 |
|  |  | *Sorghum bicolor* | 1002 (1) | 921 | 256 | 93 |
|  |  | *Setaria viridis* | 1462 (4) | 1351 | 383 | 205 |
|  |  | *Streptochaeta angustifolia* | 10 (1) | 3 | 1 | 0 |
|  | Bromeliaceae | *Ananas comosus* | 3 (1) | 2 | 89 | 0 |
| Arecales | Arecaceae | *Elaeis guineensis* | 7 (2) | 1 | 4 | 0 |
|  |  | *Phoenix dactylifera* | 6 (0) | 0 | 21 | 0 |
| Zingiberales | Musaceae | *Musa acuminata* | 13 (1) | 3 | 6 | 1 |
| Asparagales | Orchidaceae | *Phalaenopsis equestris* | 7 (1) | 0 | 0 | 0 |
|  | Asparagaceae | *Asparagus officinalis* | 31 (1) | 10 | 54 | 2 |
| Allismatales | Araceae | *Spirodela polyrhiza* | 0 | 0 | 0 | 0 |
|  |  | *Lemna gibba* | 0 | 0 | 0 | 0 |
|  | Zosteraceae | *Zostera marina* | 0 | 0 | 21 | 0 |
| Amborellales | Amborellaceae | *Amborella trichopoda* | 10 (1) | 2 | 1 | 0 |

Finally, since the number of 21-*PHAS* loci is quite high in the grasses, we asked whether these loci might be important to have on all chromosomes, for some undescribed role in chromosome biology (e.g. chromosome pairing). We analyzed the chromosomal distribution of 21-*PHAS* loci for five grasses which we had high quality genomes (*Brachypodium*, rice, *Sorghum*, maize, and *Setaria*). This showed an enrichment for 21-*PHAS* loci in some chromosomes, and a presence of at least one locus on all chromosomes (Supplemental Figure S5). While this is consistent with a hypothesis in which these loci are important for some sort of *cis* activity within the chromosome, no correlation was observed between the size of the chromosome and the count of its 21-*PHAS* loci (Supplemental Figure S5). However, this hypothetical activity would likely be limited to the grasses, given that non-grass monocots had few loci.

### miR2275 and 24-*PHAS* loci

The only known function of miR2275 is to trigger biogenesis of 24-nt reproductive phasiRNAs. We detected this miRNA in *Amborella*, and found it in both vegetative and inflorescence tissues (Figure 7). *Acorus*, a basal monocot, and *Nymphaea* a basal angiosperm, had low abundances of miR2275 in the inflorescence tissues. The early diverging grasses, *Pharus* and *Streptochaeta*, also showed moderate abundances of miR2275 in the inflorescence tissues. The highest abundances of miR2275 were in grass inflorescences (Figure 7), while in vegetative tissues the abundance was highest in *Amborella*, *Phalaenopsis*, and bamboo. The high abundance of miR2275 in grasses may reflect an increased utilization of the pathway generating 24-nt reproductive phasiRNAs.

**Figure 7.**
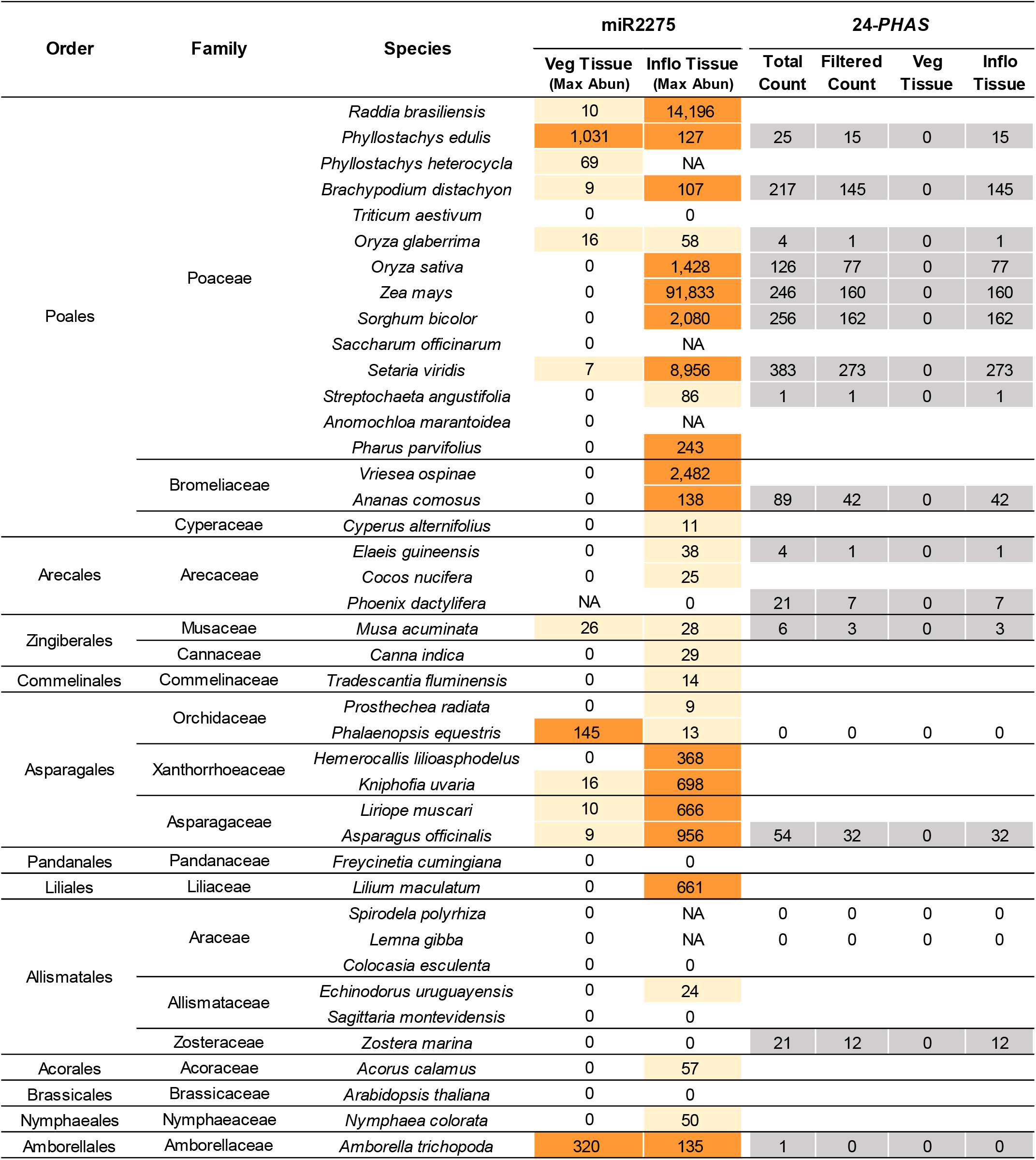
miR2275 abundance and 24-*PHAS* counts in monocots. The miR2275 maximum abundance is shown for each monocot tissue and species in which it was identified. The total and filtered 24-PHAS counts is shown for each monocot tissue and species in which it was identified. The filtered 24-*PHAS* counts for vegetative (Veg) and inflorescence (Inflo) tissues are also shown. Color legend for miR2275 abundance: beige < 100 reads, orange ≥ 100 reads. NA= not available.

We next identified 24-*PHAS* loci for 15 species for which a genome sequence was available. The highest count of 24-*PHAS* loci was in the grasses (Figure 7), although only a single 24-*PHAS* locus was identified for the early-diverged grass *Streptochaeta*. Outside the grasses, pineapple and asparagus had the highest counts of 24-*PHAS* loci, with numbers of loci that were similar to maize. We found no homologs of miR2275 in the sea grass *Zostera* or date palm, although we identified 24-*PHAS* loci in both species (12 and seven loci, respectively), perhaps consistent with asparagus or Solanaceous species (see below). We analyzed the chromosomal distribution of 24-*PHAS* loci in five grasses (*Brachypodium*, rice, *Sorghum*, maize, and *Setaria*), and as with 21-*PHAS* loci, 24-*PHAS* loci were found on all chromosomes, with counts elevated on some chromosomes (Supplemental Figure S5). Again, we observed no correlation between the size of the chromosome and the number of 24-*PHAS* loci.

We predicted the triggers of 24-*PHAS* loci (Table 1). miR2275 is the trigger for the majority of 24-*PHAS* loci in the grasses. miR2275 was also generally not found as the trigger for 24-*PHAS* loci outside grasses. For example, in *Zostera*, 21 of the 24-*PHAS* loci show no evidence that miR2275 is their trigger (Figure 7). This is consistent with recent observation from work in garden asparagus (Kakrana et al., 2018) and tomato (Xia et al., 2019), both of which have 24-nt reproductive *PHAS* loci that lack obvious miRNA triggers and indicating a diversity of initiation mechanisms for 24-*PHAS* loci outside of the grasses. In garden asparagus, it was shown that 24-nt phasiRNAs may be derived from inverted repeats even without miR2275, although how this initiates is unclear (Kakrana et al., 2018). A complete understanding of the diversity of biogenesis mechanisms for 24-*PHAS* loci will require detailed studies in these species.

### DCL5 divergence in the monocots

Earlier work on 24-nt reproductive phasiRNAs has identified a key role for DCL5 in their biogenesis (Zhang et al., 2018). This Dicer-like protein has not been well characterized outside of the grasses (or even within the grasses), so we sought to take advantage of sequenced monocot genomes and transcriptomes to determine when DCL5 emerged, and if it might coincide with any patterns of reproductive phasiRNA expression. We identified putative orthologs of DCL3 and DCL5 encoded in 12 monocot genomes plus *Amborella* and *Nymphaea*, extracted the predicted protein sequences, and performed a phylogenetic analysis (Figure 8). DCL5 orthologs were only present in *Dioscorea* and more recently diverged monocot lineages. We hypothesize that DCL5 evolved via a DCL3 tandem duplication event or whole genome duplication event, and that DCL5 emerged sometime before the diversification of *Dioscorea*. Intriguingly, DCL5 appears to have been lost independently in some orders, such as the Asparagales (*Asparagus officinalis* and *Dendrobium officinale*). Dicer-like sequences are typically long (~1,600 amino acids) with many small exons, and thus are prone to mis-annotation. Manual searches of the *Asparagus officinalis* genome sequence (Harkess et al., 2017), instead of relying solely on published annotations, revealed a partial Dicer-like sequence with homology to DCL5 (Supplemental Figure S9). However, the Asparagales still express 24-nt reproductive phasiRNAs triggered by miR2275 (Table 1), suggesting that DCL3 may have retained a redundant ancestral function. One explanation may be that species in the Asparagales have lost DCL5, but have evolved modified genomic substrates for DCL3 and miR2275 to still produce 24-nt reproductive phasiRNAs. In *Asparagus officinalis*, 24-nt reproductive phasiRNAs are often derived from inverted repeats, which may have evolved as a DCL5-independent mechanism to produce phasiRNAs (Kakrana et al., 2018). We conclude that monocots which emerged coincident with the duplication of DCL3 may have adapted diverse mechanisms for production of 24-nt phasiRNAs. Earlier diverged species including eudicots (Xia et al., 2019) likely utilized DCL3, while the specialized DCL5 emerged in later diverged species, including the grasses.

**Figure 8.**
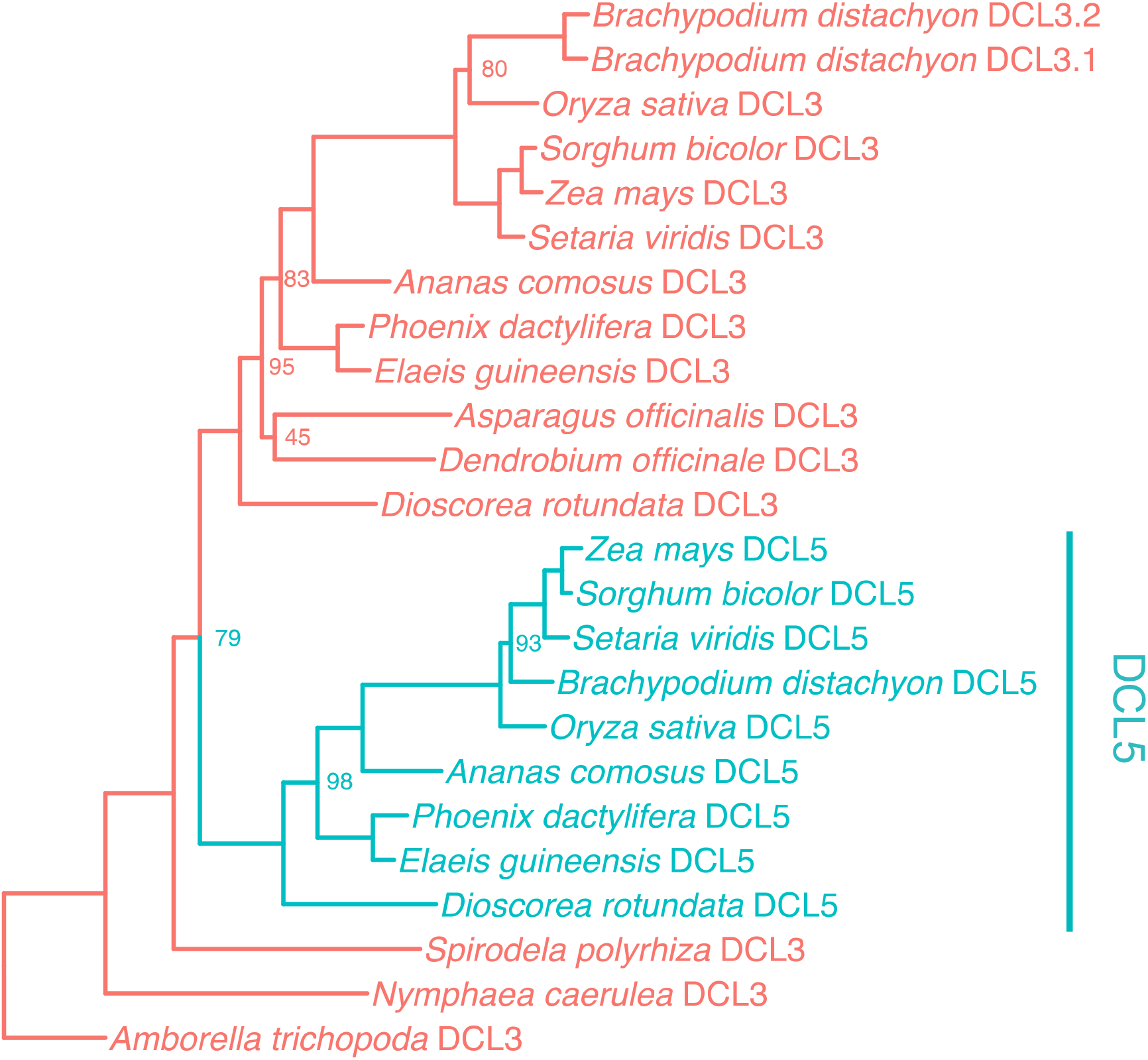
The origin of DCL5 in the monocots. A Maximum Likelihood (ML) RAxML tree of monocot-wide Dicer-like 3 (DCL3) and Dicer-like 5 (DCL5) protein sequences. Only bootstrap values less than 100 are presented on nodes.

## Discussion

Our understanding of plant small RNAs is largely derived from work focused on species with sequenced genomes, including Arabidopsis, rice, soybean, Medicago, etc. A small number of studies have surveyed more diverse species, often lacking genomic data, such as lycophytes, ferns, and diverse angiosperms (Montes et al., 2014; You et al., 2017). We focused on a poorly-sampled but diverse group of angiosperms, the monocots. We analyzed 41 species, including 38 monocot species, ranging from *Acorales* and *Arecales* to *Zingiberales*, totaling 308 sRNA libraries, including 200 sRNA libraries that were newly generated. Some observations were not surprising: the predominant size classes, 21 and 24 nucleotides, are typical of plants. Yet by calculating the ratio of 24- to 21 nt sRNAs, we found a higher ratio in the grasses and in the inflorescence tissues compared to non-grasses and vegetative tissues, respectively. We observed in the inflorescence libraries a disproportionately high level of 22-nt sRNAs compared to vegetative tissues for most grasses (*Setaria*, *Sorghum*, etc.) and for several non-grass monocots (*Tradescantia*, *Phalaenopsis*, *Zostera*, among others). There are both similarities and differences with maize; we demonstrated the significant presence of 22-nt siRNAs outside of maize, but also found that these 22-nt siRNAs have a distinct sequence composition relative to maize (Patel et al., 2018). The biogenesis and functions of these 22-nt siRNAs remains unclear. Perhaps in these species, DCL2 has a role in silencing endogenous elements, as it is the primary Dicer-like protein that produces 22 nt sRNAs, at least in Arabidopsis (Blevins et al., 2006).

Based on sequence homology, we characterized 37 miRNA families (21 highly and 16 intermediately conserved) and observed miRNA conservation patterns such as lineage-specific loss (for example, miR1507), recent evolutionary emergence (for example, miR444, miR530, and miR1432), functional diversification (absence of miR482 and emergence of miR2118 in the grasses), and emergence prior to monocots (presence of miR482, miR1432, and miR2275 in *Amborella*). Our characterization of conserved miRNAs using single-nucleotide miRNA sequence profiles revealed a position-specific nucleotide biases (at the 8th, 9th, and 19th positions) of conserved miRNA variants, potentially influencing AGO sorting. Lastly, we did not identify any significant pattern of novel, previously unannotated miRNAs that are conserved across all monocots.

Our analysis of miRNA sequence characteristics identified a number of issues with miRBase-annotated miRNAs. The analysis of sequence profiles among conserved miRNAs identified numerous strong indicators of true miRNAs. This includes the well-described 5’ and 3’ nucleotides, but there are internal nucleotides that may be important, with previously-unrecognized characteristics such as the peak of G that we observed at the 8th and 9th positions, and the peak of C at the 19th position (with a depletion of G and U). We also confirmed prior observations that misannotations persist in miRBase and may cause recurring annotation problems, such as the set of tRNA fragments represented by miR6478 and miR894/miR8155/miR8175. There is perhaps a need to track not just true annotations of miRNAs but also to track false annotations along with the explanation of why these sequences are deprecated. Our work supports the case for making community-driven improvements to miRBase (Axtell and Meyers, 2018). A related improvement would be an automated interface to provide rapid quality assessment of miRNAs, assign unique miRNA identifiers, and track targets including whether or not the loci yield phasiRNAs.

Finally, we showed that the miR2118 and miR2275 triggers of reproductive phasiRNAs, and their associated genomic *PHAS* loci, are prevalent in angiosperms. This overall observation is consistent with recent work on eudicots (Xia et al., 2019). Moreover, we demonstrated conservation of miR2118 and miR2275 across monocot species, with evidence of expression in the inflorescence tissues of all grasses, as well as limited expression in some vegetative tissues in a few non-grasses. Similarly, we concluded that 21- and 24-*PHAS* loci (the sources of reproductive phasiRNAs) are particularly numerous and abundantly expressed in inflorescence tissues of the grasses. *In silico* trigger identification in the grasses determined that miR2118, miR390, and miR2275 are triggers of most 21-*PHAS* loci, the handful of *TAS3* loci, and 24-*PHAS* loci, respectively. Prior work in species outside of the grasses has also demonstrated that 24-nt reproductive phasiRNAs may be generated by non-canonical pathways (Kakrana et al., 2018; Xia et al., 2019). In fact, this variation in the utilization of miR2275 may reflect an evolutionary period of divergence in 24-nt reproductive phasiRNA biogenesis, coincident with changes occurring in the divergence of DCL3 and DCL5. We narrowed the phylogenetic placement in monocot evolution during which DCL5 likely emerged. In the lineages that diversified following the evolution of DCL5, there is presence/absence variation for DCL5, suggesting that neofunctionalization was incomplete and its necessary function in reproductive phasiRNA biogenesis observed in grasses (Song et al., 2012) was not yet fixed. Future functional studies should test the specialization and activity of DCL5 from the genomes of these first lineages to inherit it.

In conclusion, we are in a period of rapid, large-scale data acquisition, including both genomes and the transcript data from these genomes. The interpretation of these data requires increasingly sophisticated methods amenable to high-through analyses. One particularly intriguing branch of research focuses on machine learning applications, which in our experience is transforming small RNA informatics (Patel et al., 2018). Future applications of machine learning may provide even deeper insights into comparative analysis, elucidating uncharacterized aspects of small RNA sequences, and elevating our understanding of miRNAs, phasiRNAs, and other classes of plant RNAs.

## Methods

### Plant material

Plant materials were collected from various locations and are detailed in Supplemental Table 1. Plant tissues were dissected manually and when necessary, using a stereomicroscope for magnification and a 2-mm stage micrometer (Wards Science, cat. #949910). Tissues were flash frozen in liquid nitrogen and stored at −80°C until RNA extraction was performed.

### RNA extraction

Samples were ground in cold mortars and pestles using liquid nitrogen. Total RNA isolation was performed using the Plant RNA Reagent (Thermo Fisher, cat. no. 12322012). For anther and other tissues with limited amount of plant material, we used the detailed protocol for RNA extraction previously published (Mathioni et al., 2017).

### Small RNA size selection, library preparation and sequencing

Small RNA size selection and library preparation using the TruSeq Small RNA Library Prep Kit (RS- 200-0012, Illumina) were performed following the detailed protocol previously published (Mathioni et al., 2017). All small RNA libraries were sequenced using single-end mode with 51-bp on an Illumina HiSeq 2500 Instrument in the University of Delaware Sequencing and Genotyping Center at the Delaware Biotechnology Institute.

### PARE library preparation and sequencing

PARE libraries were constructed as described previously (Zhai et al., 2014), with modifications as follows: the amount of total RNA used as starting material was 10 to 20 ug, depending on the availability of each sample; the incubation time for the 5’ adapter ligation was 2 hours; the incubation time for the reverse transcription was 2 hours; the second strand cDNA synthesis was performed with 10 cycles instead of 7 cycles; the incubation time for the MmeI digestion was 2 hours; the incubation time of the 3’ double-strand DNA adapter ligation was performed overnight (~ 8 hours); the final PCR amplification of the PARE library was performed with 18 cycles instead of 15 cycles. These changes were necessary because the total RNA starting amount was much lower than the amount recommended in the original protocol (40 to 75 ug). All the other parameters were kept as described in the original protocol. All PARE libraries were sequenced using single-end mode with a 51 nt read length on an Illumina HiSeq 2500 Instrument in the University of Delaware Sequencing and Genotyping Center at the Delaware Biotechnology Institute.

### Small RNA and PARE data processing

The small RNA data were processed as previously described (Mathioni et al., 2017). The PARE data were processed using sPARTA as previously described (Kakrana et al., 2014) with updates available at https://github.com/atulkakrana.

### MicroRNA analysis

We used two strategies to analyze the microRNAs. First, we generated a unique list of miRBase entries from version 21, with all mature miRNA sequences from the available plant species. We used this list to query our small RNA dataset for mature miRNA sequences. The criteria used for the query were based on sequence similarity (see below). In the second strategy, we ran a *de novo* miRNA prediction using both a new prediction package called *miRador* (https://github.com/rkweku/miRador) and using ShortStack (Johnson et al. 2016) for the each of species with genome sequences available.

### Homology based identification of miRNA sequences in diverse monocots using BLAST

Due to a lack of sequenced genomes in monocots, we conducted a homology-based search to identify miRNAs in 40 monocot species. We added *Arabidopsis thaliana* (Arabidopsis) as a control, making it a list of 41 species. We merged all sRNA libraries from these 41 species into one and built a BLAST database (BLASTdb) (Altschul et al., 1990). The command line used was makeblastdb -in file -out name -dbtype nucl -title title. Each sequence name in this file embeds information about the species name, raw read count, and tissue (vegetative or inflorescence). To compare our sRNA sequences in the BLASTdb, we collapsed a Viridiplantae-specific, non redundant, mature miRNA sequences from miRBase version 21 (Kozomara and Griffiths-Jones, 2014). This set of unique miRNA sequences completed our reference set. To identify miRNAs present in our BLASTdb, we aligned the BLASTdb to the reference set using BLASTN. The command line used was blastn -query file -strand plus -task blastn-short -db name -out file - perc_identity 75 -num_alignments 200 -no_greedy -ungapped -outfmt “6 qseqid query sseqid pident length qlen qstart qend slen sstart send gaps mismatch positive evalue bitscore”. BLASTN was performed with –ungapped and -no_greedy to facilitate end-to-end and non-greedy alignment, respectively. Here, we utilized “blastn-short”, which is optimized for sequences less than 30 nucleotides. We used 75% percent identity for BLASTN sequence scan to account for ≤ 4 mismatches between a mature miRNA from the reference set and its homolog (subject sRNA sequence) in the BLASTdb. Output file contained fields identified as qseqid, query, sseqid, pident, length, qlen, qstart, qend, slen, sstart, send, gaps, mismatch, positive evalue, and bitscore in a TAB separated format.

To process miRNA annotation results from BLAST, the output from BLAST was filtered to determine the valid homologs. We used a custom python script for this filtration process. The filtration process was as follows: (1) we processed the tabular results from BLAST to compute 5’- and 3’-end overhangs, mismatches, matches, and total variance. Total variance was a sum of the nucleotides that were not aligned, including no more than two nucleotides on 5’- and 3’-end overhangs and mismatches (≤ 4). The total variance cutoff was set to 5. Subject sequences (aka candidate homologs) satisfying this cutoff were given a status of “pass”, otherwise “fail”. In addition to the output from BLAST, we added extra columns (hang5, hang3, match, mismatch, unalign, totalvariance, and status). (2) We only kept the subject sRNA sequences which are of length between 20-22 nt. (3) When a subject sRNA sequence matched two or more mature miRNAs from the reference set, the best match was determined as the alignment that contained the highest bitscore. (4) For each sRNA subject sequence that passed these criteria, we determined the raw read count in the inflorescence and in the vegetative tissues across all sRNA libraries used. All of the potential homolog candidates were chosen based on their raw read counts more than 99 reads in either vegetative or inflorescence tissues. We used another custom python pipeline to obtain these read counts and added three extra columns (Homologous Sequence, Vegetative Raw Read Count and Reproductive Raw Read Count) to yield the final results.

In order to obtain the count of miRNA families, we used three-letter miRBase (v21) codes (i.e. identifiers lower than miR1000) and generated the set of Viridiplantae-specific miRNA families using a custom python script. The same script searched the list of entries from the final results and collapsed these candidate sequences using a column query into the list of miRNA families, demonstrating the conservation of families of miRNAs in these monocot species.

### *De novo* miRNA prediction for novel miRNAs

sRNA libraries were trimmed (Patel et al., 2015) and mapped to their respective genomes using Bowtie (Langmead et al., 2009). miRNAs were then predicted in all libraries utilizing a version of miREAP software (https://sourceforge.net/projects/mireap/). These miRNAs were assessed for their similarity to known miRNAs using BLAST. Predicted miRNAs that had five or fewer differences were then classified as members of those miRNA families.

### PhasiRNA analysis

Phased small interfering RNA (*PHAS*)-generating loci were identified using the *PHASIS* pipeline (https://github.com/atulkakrana/PHASIS) (Kakrana et al., 2017). Triggers for these *PHAS* loci were further identified using the *phastrings*, a component of the *PHASIS* pipeline.

### Dicer-like gene family and phylogeny

*De novo* gene families were circumscribed using a set of diverse monocot genomes from Phytozome v12.1 (*Ananas comosus*, *Asparagus officinalis, Brachypodium distachyon, Musa acuminata, Oryza sativa, Sorghum bicolor, Zostera marina, Setaria viridis,* and *Zea mays*), plus *Amborella trichopoda)* using OrthoFinder (v2.2.1) (Emms and Kelly, 2015). DCL3 and DCL5 proteins were contained in a single orthogroup. Additional monocot sequences were added from manual BLASTP searches of the *Dioscorea rotundata* genome (Tamiru et al., 2017), and from NCBI for *Elaeis guineensis*, *Phoenix dactylifera*, and *Dendrobium caternatum*.

Only proteins that were full, canonical Dicer-like proteins (including a helicase domain, Dicer domain, PAZ, and two tandem RNAse III domains) were retained, except in a single case of possibly misannotated recent DCL3 paralogs in *Brachypodium*. This led to the exclusion of several Dicer-like genes in the analysis, which may be pseudogenes, incorrectly assembled or unannotated genomic regions, or non-canonical Dicer-like proteins that perform a currently unknown function (Supplemental Figure S9).

Complete proteins were aligned using default settings in PASTA (v1.6.4) (Mirarab et al., 2015), followed by a Maximum Likelihood (ML) gene tree using RAxML (v8.2.11) (Stamatakis, 2014) over 100 rapid bootstraps with options “-x 12345 –f a –p 13423 –m PROTGAMMAAUTO”. Trees were visualized and manipulated in ggtree (Yu et al., 2017).

## Accession Numbers

The accession numbers for all the small RNA libraries generated in this study and all the libraries from published studies and used in this study for comparison purposes are listed in Supplemental Table S2.

## Author Contributions

B.C.M. designed the study; S.M.M. and S.A. collected plant material and dissected plant tissues; S.M.M. and S.A. prepared small RNA libraries; P.P. and A.K. processed the data; S.M.M., P.P., A.E.H., and A.K. analyzed the data; P.P., S.M.M., A.E.H. and B.C.M wrote the manuscript. All authors read and approved the manuscript.

## Acknowledgements

We would like to thank Robert J. Orth and Andrew J. Johnson for providing *Zostera marina* samples; Elizabeth Kellogg for providing *Streptochaeta angustifolia* and *Pharus parvifolius* samples; Malia Gehan for providing *Setaria viridis* seeds; Nadia Shakoor for providing *Sorghum bicolor* seeds; the Longwood Gardens (https://longwoodgardens.org/) for providing samples from *Vriesea ospinae*, *Cyperus alternifolius*, *Canna indica*, *Tradescantia fluminensis*, *Prosthechea radiata*, *Freycinetia cumingiana*, *Echinodorus uruguayensis*, *Sagittaria montevidensis*, and *Nymphaea colorata*; the Chanticleer Garden (http://www.chanticleergarden.org/) for providing *Acorus calamus* samples; the T.S. Smith and Son’s Farm (Bridgeville, DE) for providing *Asparagus officinalis* samples. We thank Deepti Ramachandruni and Mayumi Nakano for assistance in data handling and submission, as well as members of the Meyers lab for their helpful discussions. We would like to thank Brewster (Bruce) Kingham, Olga Shevchenko, and Summer Thompson in the University of Delaware Sequencing and Genotyping Facility. This work was supported by funding provided by the National Science Foundation awards # 1339229 to B.C.M and #1611853 to A.E.H, and resources provided by the Donald Danforth Plant Science Center.

---

**Supplemental Figure S1.**
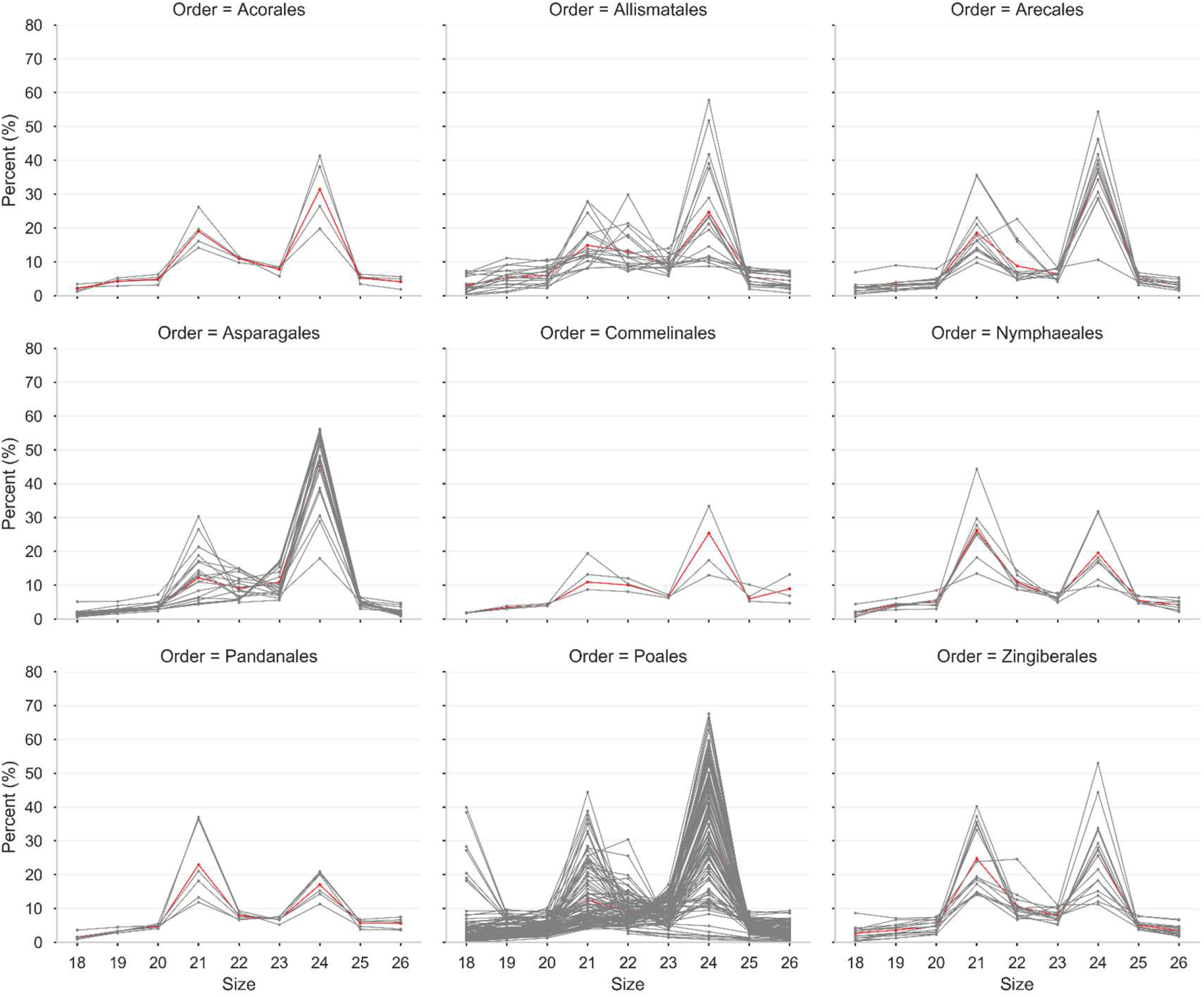
Size distribution of sRNA sequences in nine different plant orders used in this study. The relative proportion of sRNAs are displayed as percentages (Y-axis) for each size category (X-axis) in total reads in 200 individual sequenced libraries, denoted by gray lines, across 28 plant species in 9 different plant orders. Reads longer than 26 nt are not included in this study. Red line indicates average proportions of sRNAs in all libraries in that plant order.

**Supplemental Figure S2.**
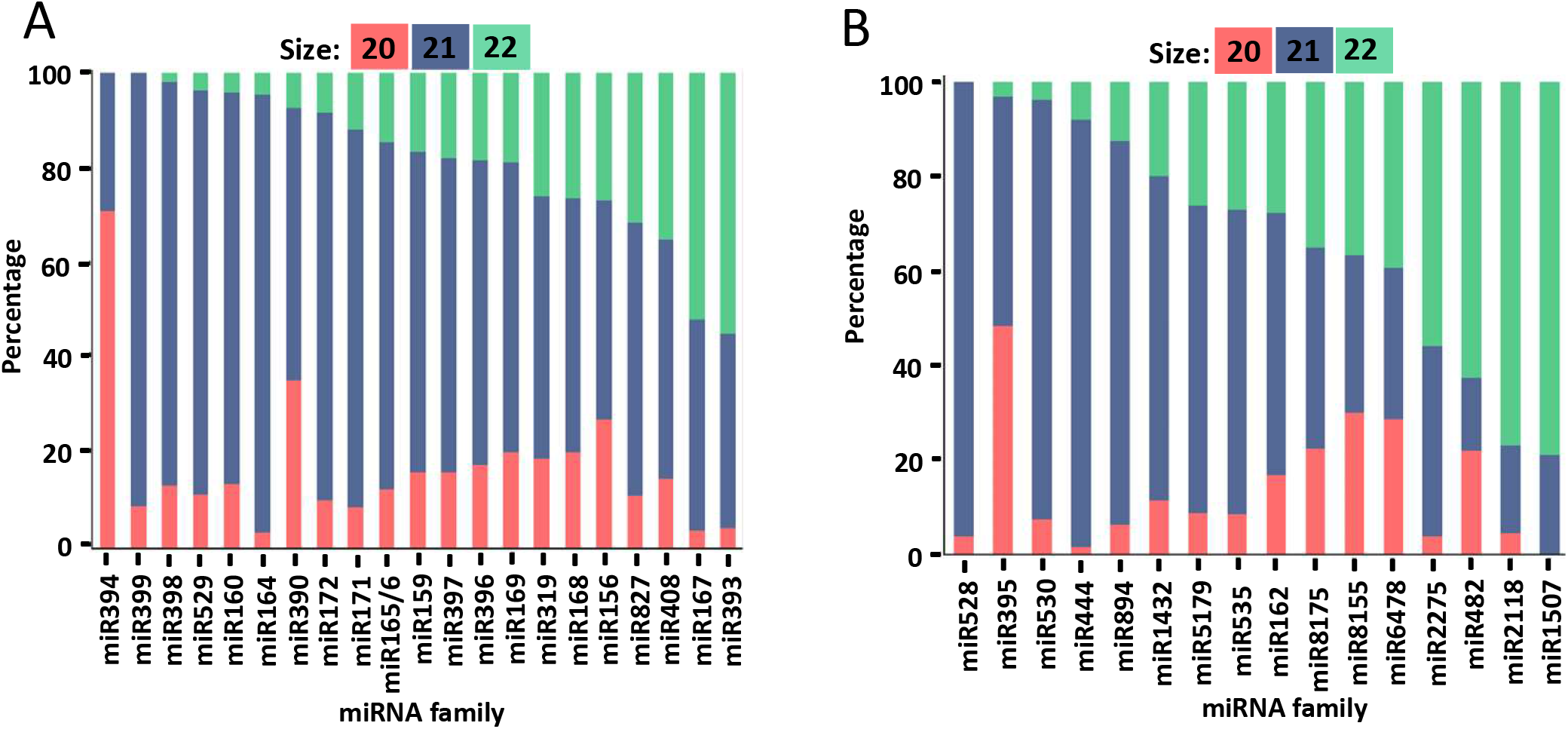
Size distribution of conserved miRNA variants by size. (A-B) The stacked bar plots show the size distribution of reads as a percentage (Y-axis, denoted 20-mers in red; 21-mers in blue; 22-mers in green) sizes between 0 and 100 in the conserved miRNA families. Similar to figure 3, conservation is defined by miRNA families identified in more than 10 species out of all 41 species examined. Bar plots are sorted from low to high percentage of 22-mers. (A) Size distribution in the most conserved 21 miRNA families (X-axis). (B) Size distribution in the intermediate conserved 16 miRNA families.

**Supplemental Figure S3.**
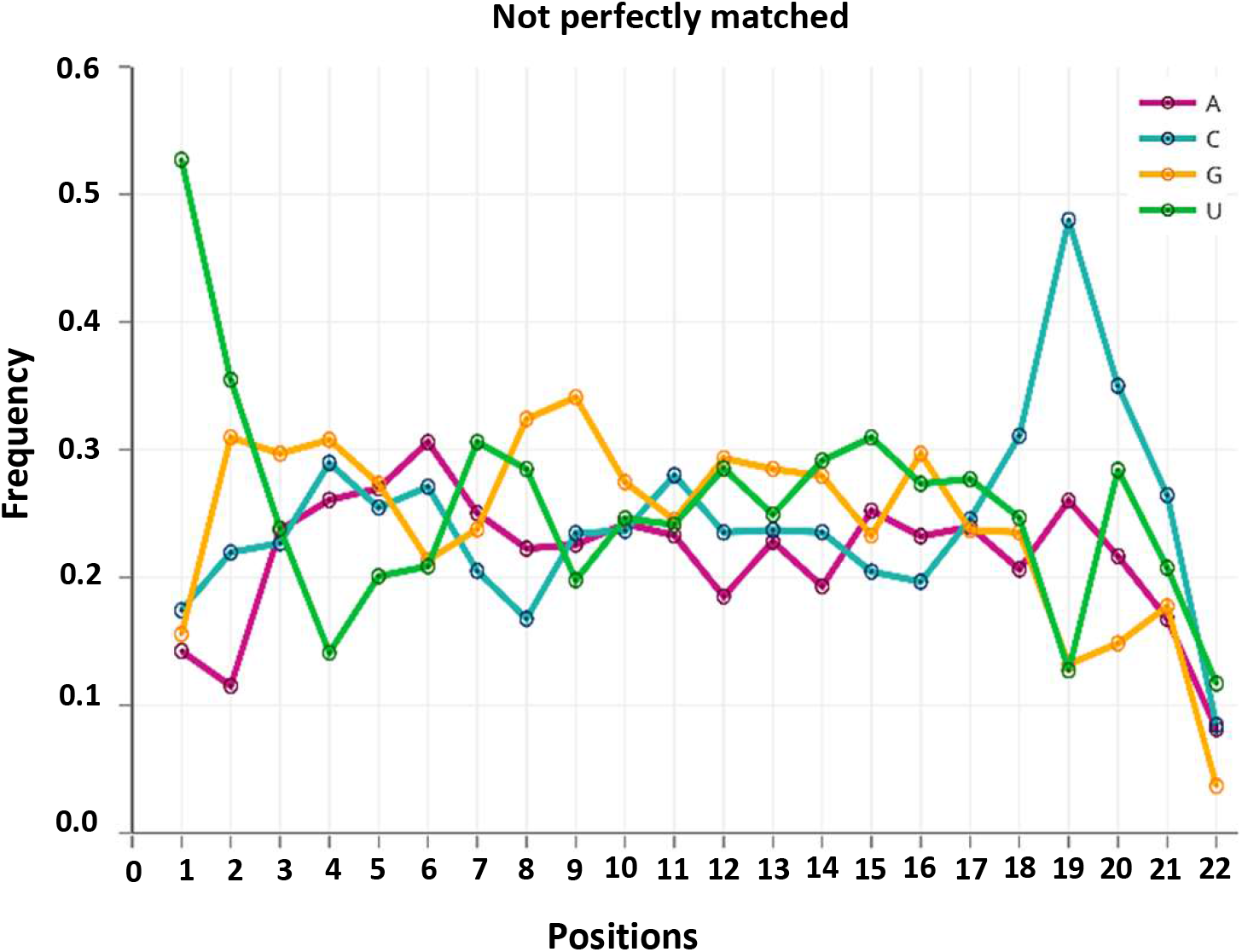
Single-nucleotide sequence profiles of unique candidate miRNAs. The unique candidate miRNAs were from Figure 4, panel C, those not perfectly matching (n= 1657) to the miRBase miRNAs.

**Supplemental Figure S4.**
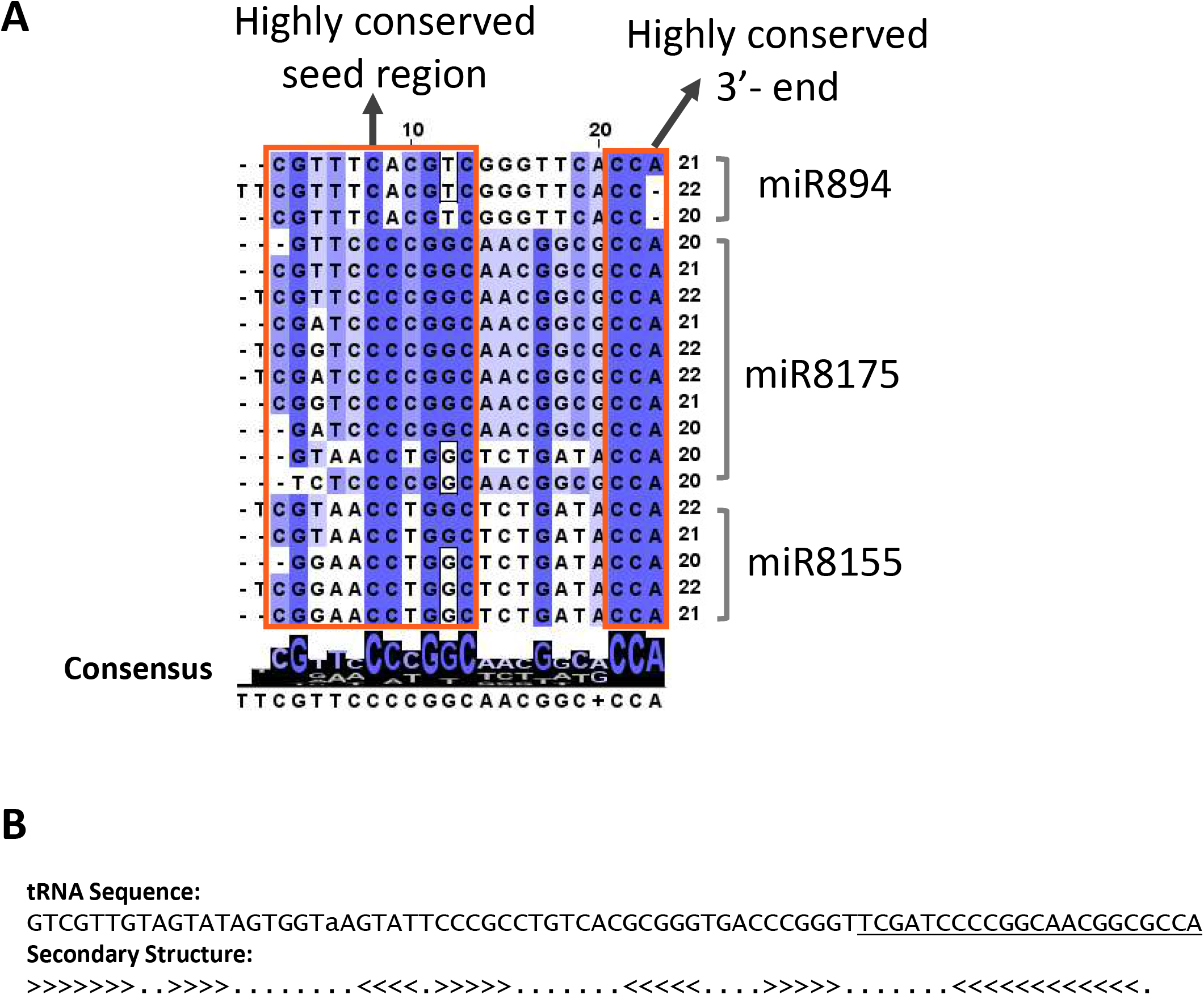
Sequence analysis of homologs miR894, miR8155, and miR8175 identifies tRNA homology. (A) The degree of sequence conservation along the miRNAs is represented by the color, with a dark color denoting a high level of conservation and a light color denoting a low level for each nucleotide. The consensus sequence of the alignment is displayed below with sequence logos. (B) Sequence of the Arabidopsis aspartyl-tRNA synthetase from: http://plantrna.ibmp.cnrs.fr/plantrna/trna/9988.

**Supplemental Figure S5.**
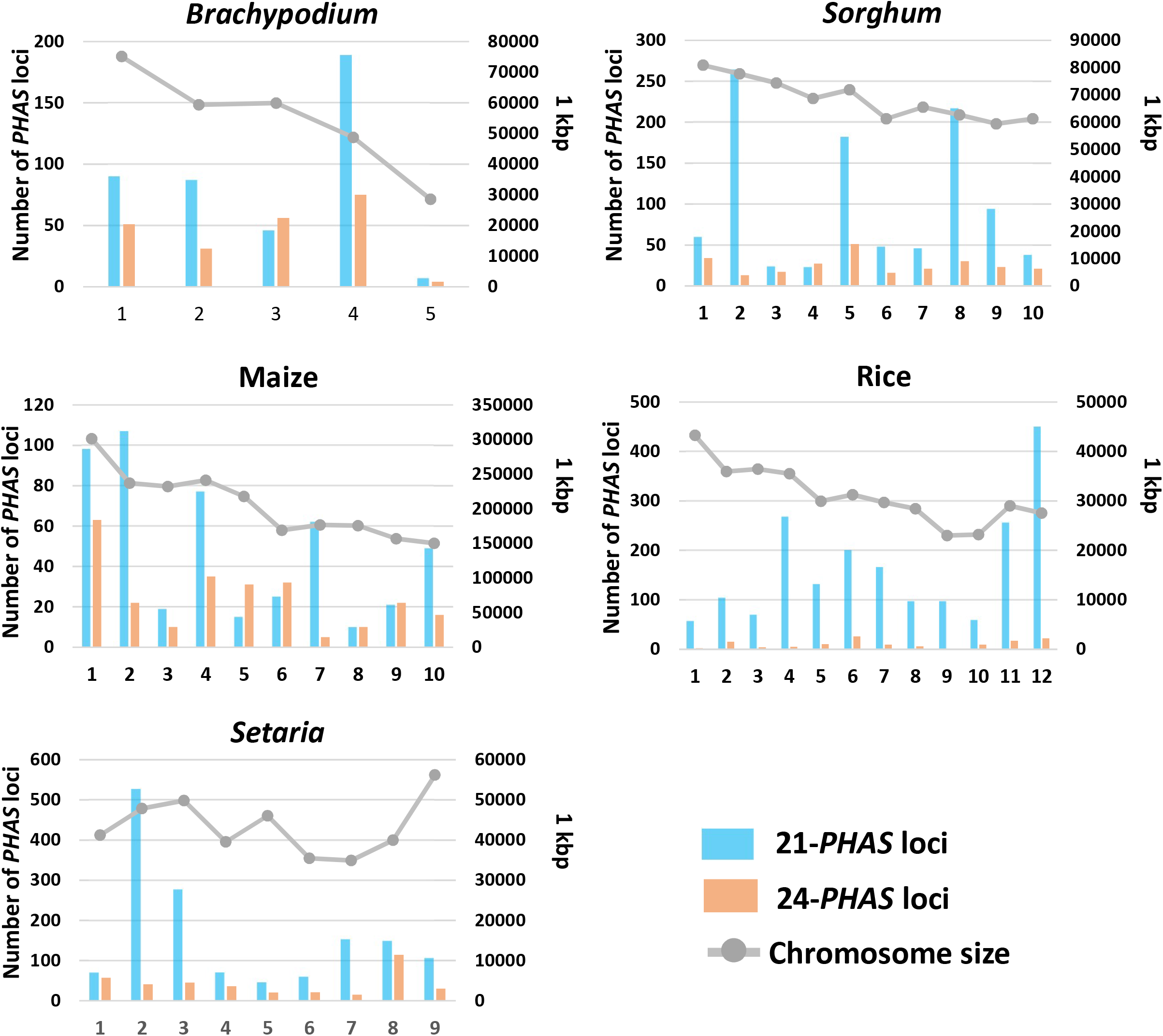
Count of 21-and 24-*PHAS* loci per chromosome in five grass species. The left Y-axis shows the counts of 21-(blue) and 24 - (orange) PHAS loci, and X-axis shows the chromosome for each of five grass genomes (as indicated). Line plots in grey indicate the size of chromosomes, denoted on a secondary Y-axis (at right) in 1 kbp units.

**Supplemental Figure S6.**
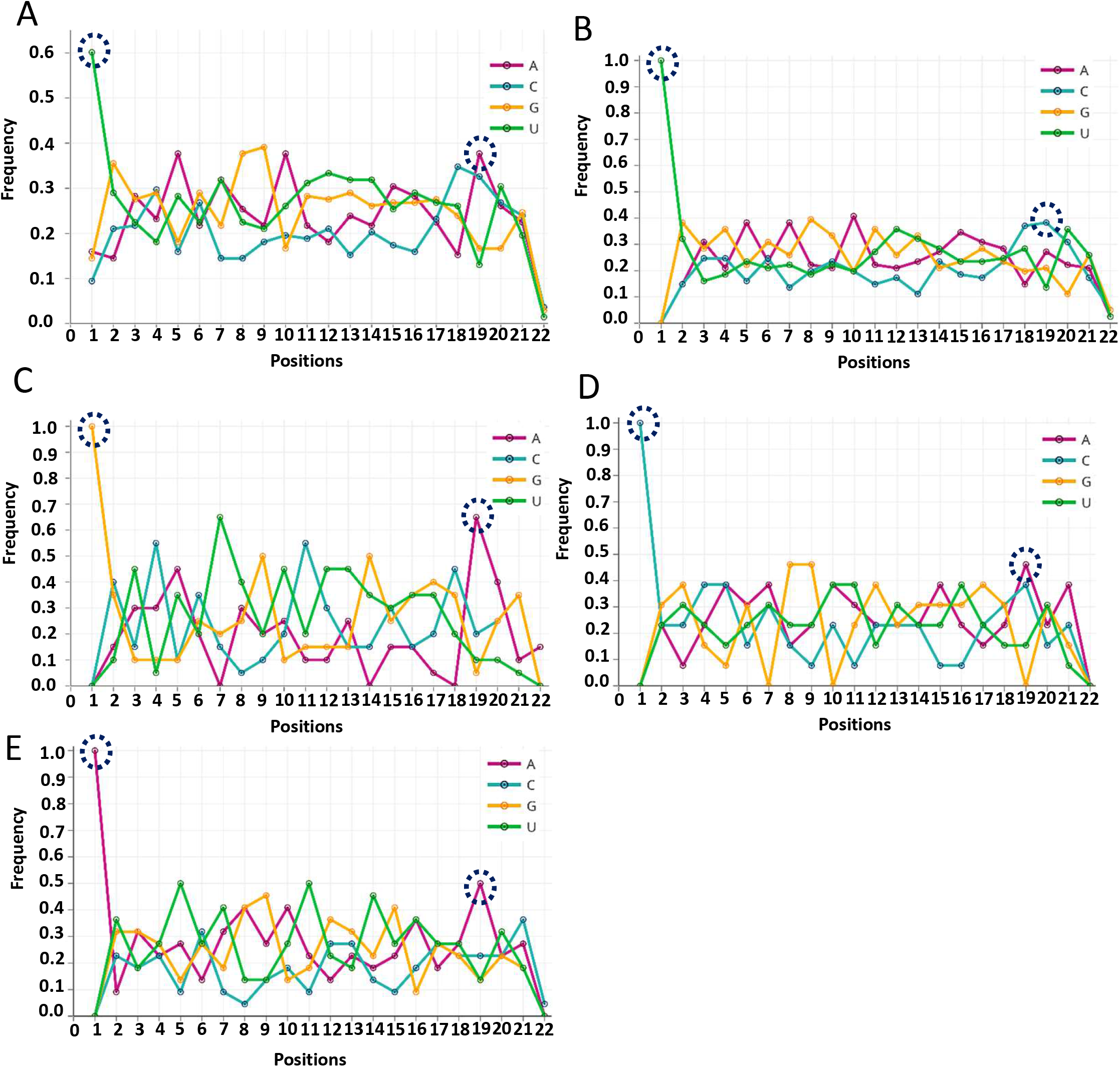
Arabidopsis 5’ U miRNAs share a distinctive C in the 19th position with high-confidence monocot miRNAs. (A) Single-nucleotide sequence profiles of Arabidopsis miRNAs (n=138) from miRBase version 21, filtered based on abundance characteristics, as described in the main text. The plot is as described in Figure 4, with the nucleotides of interest (5’ end and 19^th^ position) circled. (B to E) Four subsets of miRNAs from panel A were analyzed separately for their single-nucleotide sequence profile, based on their 5’ nucleotide composition.

**Supplemental Figure S7.**
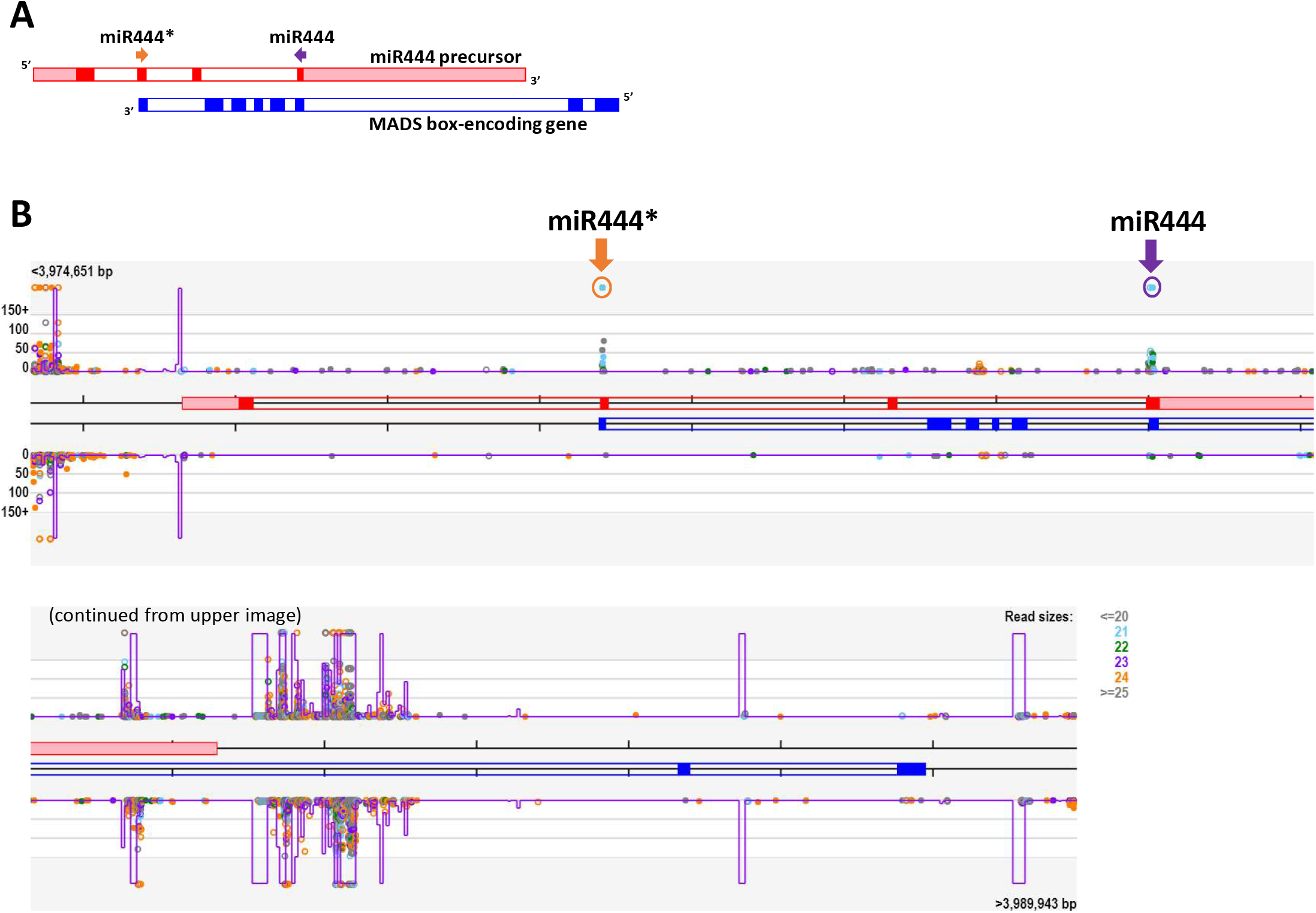
The genomic structure of the natural antisense microRNA miR444 is conserved in pineapple. A. Genomic arrangement of the natural antisense microRNA (“nat-miRNA”) miR444 as described by Lu et al. (2008). One strand generates the miRNA precursor, while the complementary strand encodes a MADS-box; transcripts overlap in the region yielding the miRNA and the miRNA target (as well as the miRNA*-generating region), while other exons are intronic or upstream relative to the complementary strand. B. Screenshot from the Meyers lab genome browser of the locus Aco003017 (on upper strand) and the complementary bottom strand, in pineapple. Purple and orange arrows and circles indicate miR444 and miR444* sites, respectively. Light blue dots indicate 21-nt reads mapped to the locus; other dots as indicated in the key at lower right, with the position on the y-axis indicating read abundance in TP20M. Four exons of antisense strand (genes on top strand) are shown in the orientation from 5’ to 3’ as denoted by red or pink color. Eight exons on the sense strand of the gene encoding the MADS box protein (on bottom strand) are denoted by the solid blue color. Purple lines within the screenshot are a k-mer frequency, indicative of repetitiveness of the region.

**Supplemental Figure S8.**
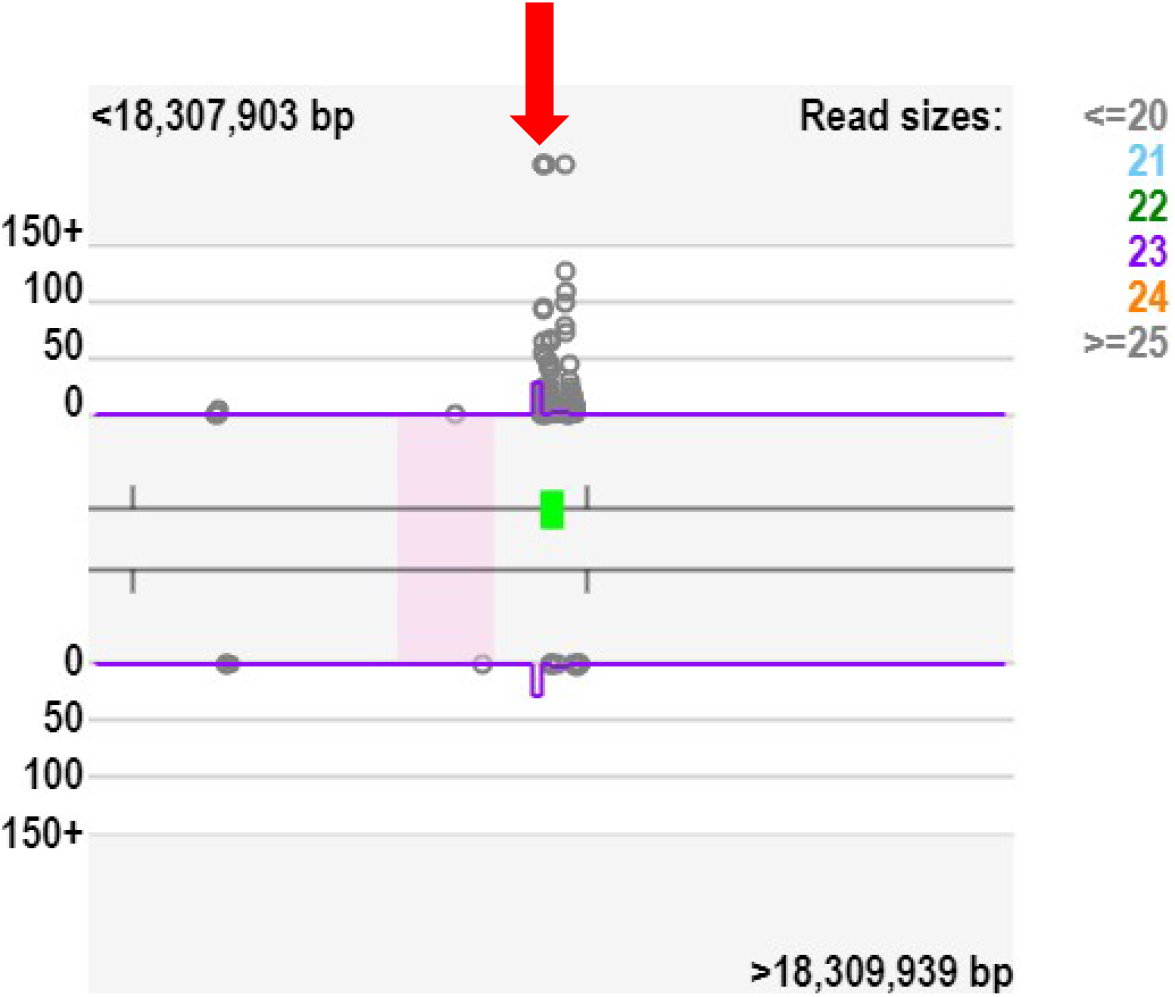
miR6478 is derived from a tRNA precursor in Arabidopsis. Screenshot from our genome browser of a locus (AT1G49460) in Arabidopsis. The red arrow shows where the annotated miR6478 sequence maps on the locus and green box indicates the annotated tRNA precursor. Nearly all small RNAs from this locus are either less than 21 or greater than 24 nucleotides, and map to multiple locations in the Arabidopsis genome, consistent with miR6478 as a tRNA fragment rather than a true miRNA.

**Supplemental Figure S9.**
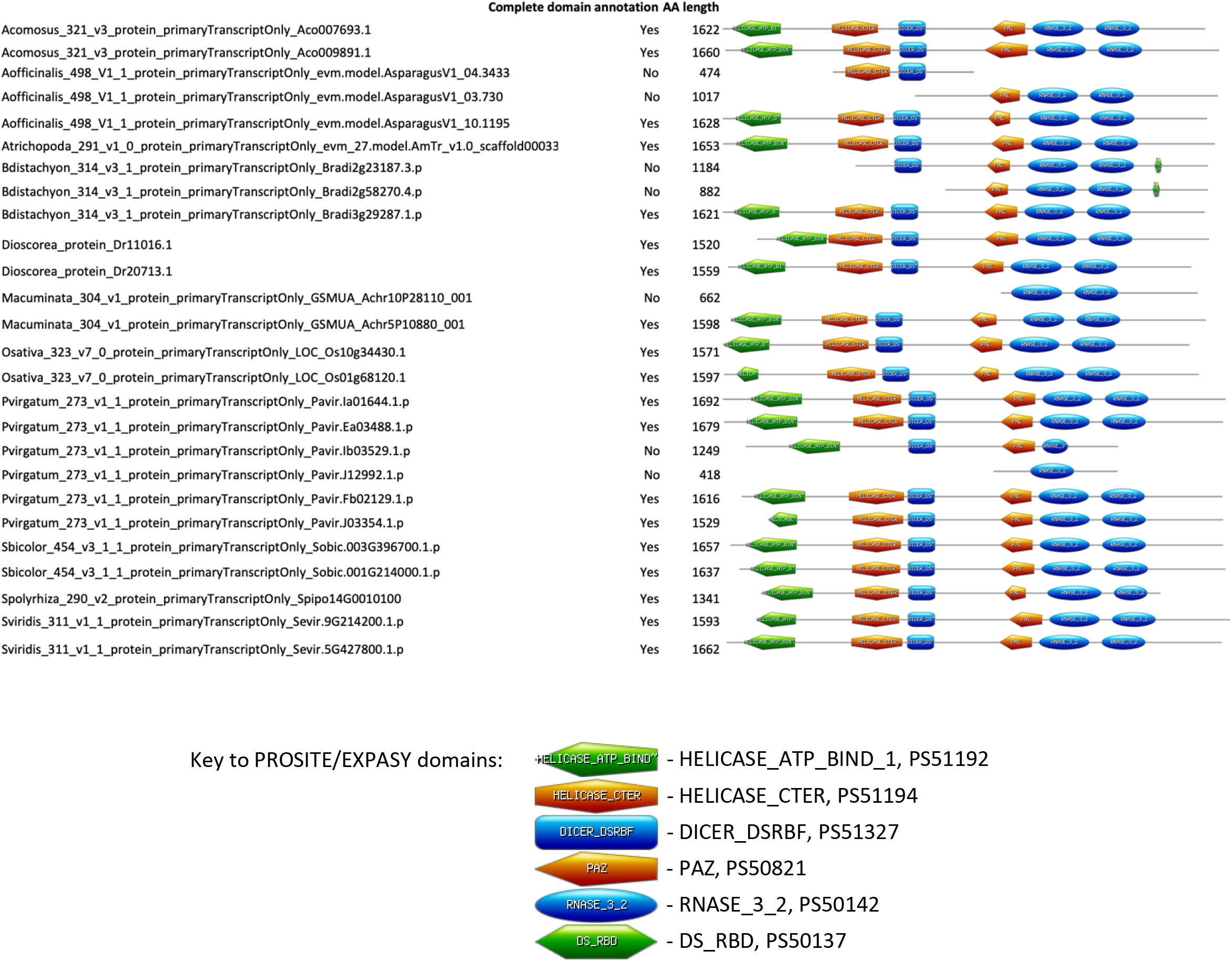
DCL3- and DCL5-like sequences and predicted domains from across the monocots based on genome annotations. For each protein sequence, domains were predicted using the ProSite server (https://prosite.expasy.org/).

